# Single Cell Spatial Analysis Reveals the Topology of Immunomodulatory Purinergic Signaling in Glioblastoma

**DOI:** 10.1101/2022.01.12.475925

**Authors:** Shannon Coy, Shu Wang, Sylwia A. Stopka, Jia-Ren Lin, Rumana Rashid, Clarence Yapp, Cecily C. Ritch, Prasidda Khadka, Michael Regan, Jaeho Hwang, Patrick Y. Wen, Pratiti Bandopadhayay, Keith L. Ligon, Nathalie Y.R. Agar, Peter K. Sorger, Mehdi Touat, Sandro Santagata

## Abstract

Glioblastoma develops an immunosuppressive microenvironment that fosters tumorigenesis and resistance to current therapeutic strategies. Here we use multiplexed tissue imaging and single-cell RNA-sequencing to characterize the composition, spatial organization, and clinical significance of extracellular purinergic signaling in glioblastoma. We show that glioblastoma exhibit strong expression of CD39 and CD73 ectoenzymes, correlating with increased adenosine levels. Microglia are the predominant source of CD39, while CD73 is principally expressed by tumor cells, particularly in tumors with amplification of *EGFR* and astrocyte-like differentiation. Spatially-resolved single-cell analyses demonstrate strong spatial correlation between tumor CD73 and microglial CD39, and that their spatial proximity is associated with poor clinical outcomes. Together, this data reveals that tumor CD73 expression correlates with tumor genotype, lineage differentiation, and functional states, and that core purine regulatory enzymes expressed by neoplastic and tumor-associated myeloid cells interact to promote a distinctive adenosine-rich signaling niche and immunosuppressive microenvironment potentially amenable to therapeutic targeting.

## INTRODUCTION

Cancers develop a wide range of molecular mechanisms to evade immune surveillance^1^. These include the expression of ‘checkpoint’ proteins that suppress cytotoxic T cells^2^, production of local and systemic inflammatory mediators^3^, recruitment of immunomodulatory immune cells^4^, and alteration of the tumor metabolic niche^5^. Together, these mechanisms are hypothesized to shape the composition and properties of the tumor microenvironment^6^. However, many unresolved questions remain regarding how immunosuppressive mechanisms are deployed in human cancer tissues, including the roles of various tumor and immune cells in generating the permissive immunologic conditions necessary for tumorigenesis and progression, and how these cells functionally and physically interact within the tumor microenvironment. There is therefore an important need to generate detailed maps that reveal the cell types and states present within human tumors and their spatial relationships.

Recently developed methods for multiplexed tissue imaging permit characterization of the heterogeneous cell states present in tumor ecosystems directly from human cancer resection tissues^7–12^. Cyclic immunofluorescence (CyCIF) is one such method, enabling quantification of up to 60 antigens at single-cell resolution. CyCIF can be used to identify multiple cell lineages and states and map their spatial organization and interactions^13^. Using such methods to generate 2D and 3D tumor atlases, such as those developed by the Human Tumor Atlas Network (HTAN)^14^, has revealed substantial spatial and molecular variation within human tumors, and population-level regulation of tumor-immune interactions^15–17^.

Glioblastoma is the most common adult primary brain tumor and a highly aggressive disease with a dismal prognosis^18^. Median survival is approximately one-year and long-term survival is extremely rare. Numerous investigational therapies including immunomodulatory approaches have failed to provide substantial clinical benefits^19^. PD-1 checkpoint pathway inhibitors initially showed little clinical efficacy in recurrent/residual glioblastoma including those with high tumor mutation burden^20–23^. However, neoadjuvant administration of PD-1 inhibitor was found to improve survival, particularly in steroid-naïve patients, though effects remain modest and longterm survival has not yet been achieved^24,25^. These results suggest that continued refinement of immunomodulatory approaches may be a promising avenue to improve survival in glioblastoma, and modulation of additional immunoregulatory pathways may provide complementary or synergistic anti-tumor effects.

Purine metabolites such as ATP and adenosine are phylogenetically ancient biochemical compounds that link genetics, metabolism, and cell behavior in diverse organisms from bacteria to humans^26^. Purines are released into the extracellular space in a wide range of biological processes including stress, apoptosis, and necrosis, and extracellular levels of purine metabolites are tightly controlled. Purines are critical mediators of inflammatory signaling in normal and pathologic tissues and modulate the function of neutrophils, macrophages, NK cells, and lymphocytes^27^. In the central nervous system, extracellular purine metabolite levels have special significance, as they may also function as neurotransmitters and neuroregulatory ligands which signal to neuronal and glial populations at both synapses and extra-synaptic sites^28^.

Extracellular ATP and ADP are hydrolyzed to AMP via the enzymatic activity of CD39 (*ENTPD1*), an integral plasma membrane ectonucleotidase expressed by immune cells and vascular endothelium. AMP is catabolized to adenosine via ecto-5’-nucleotidase CD73 (*NT5E*), a glycosyl-phosphatidylinositol (GPI)-anchored extracellular protein which is typically expressed by regulatory T cells (Tregs) and other immune cells. Conversion of AMP to adenosine is the rate-limiting step in the catabolism of extracellular purines, and CD73 is therefore a critical regulator of local purinergic signaling and inflammatory responses.

The extracellular purine pathway is an important regulator of tumorigenesis^29^. Aberrant CD73 expression occurs in numerous cancers, and is hypothesized to result in elevated adenosine levels, which may promote tumor cell proliferation, survival, and immune evasion via binding to adenosine receptors on multiple cell types^30,31^. CD73 knockout in mouse models augments antitumor immunity, increasing infiltrating T cells and reducing metastases, suggesting that CD73 inhibition may be an effective immunomodulatory therapeutic approach^32^. Prior studies have suggested that CD73 contributes to glioblastoma pathogenesis through dysregulation of purinergic signaling^33–37^, and expression of CD73 has been correlated with increased tumor cell proliferation^30^, cell adhesion^38^, invasiveness, and NK cell infiltration^39^. However, most analyses have examined non-physiologic glioma cell lines, transgenic animal models, or small sets of tumor resections, and none to date have characterized spatial and population-based features of the immune microenvironment in human glioblastoma tissue.

In this study, we use multiplexed tissue imaging, single-cell RNA-sequencing, and tissue mass spectrometry imaging to explore the activity and significance of immunomodulatory purinergic signaling in human glioblastoma. We show that CD73 upregulation correlates with tumor genotype and differentiation state, tissue adenosine levels, and poor patient outcome. Moreover, we demonstrate that spatial interactions between tumor and myeloid cells mediate purinergic metabolism and outcome in glioblastoma. These results suggest that inhibition of purine signaling represents an attractive immunotherapeutic strategy in glioblastoma and confirm that population-based analyses of multiple cell types and states and their spatial distribution in native tumor tissue may be necessary to effectively identify and target critical regulators of tumorigenesis.

## RESULTS

### Elevated CD73 levels and purine pathway activity in glioblastoma

To evaluate the hypothesis that CD73 activity and dependence vary across cancer types, we first analyzed bulk mRNA sequencing data from The Cancer Genome Atlas (TCGA; Pan-Cancer Atlas, n=10,071 samples, 32 tumor types). This showed that glioblastomas, sarcomas, and carcinomas of the thyroid and liver expressed the highest levels of *NT5E/CD73* (**Fig. 1a**). *CD73* expression has been linked with aggressive behavior and poor clinical outcomes in each of these tumors^39–41^. Genomic alterations such as mutations, chromosomal copy-number changes, and rearrangements are uncommon for *CD73* in glioblastoma (<0.5%) and other tumor types (**Supplementary Fig. S1a**), and mRNA levels were not significantly elevated in tumors with gene amplification (**Supplementary Fig. S1b**).

**Figure 1:**
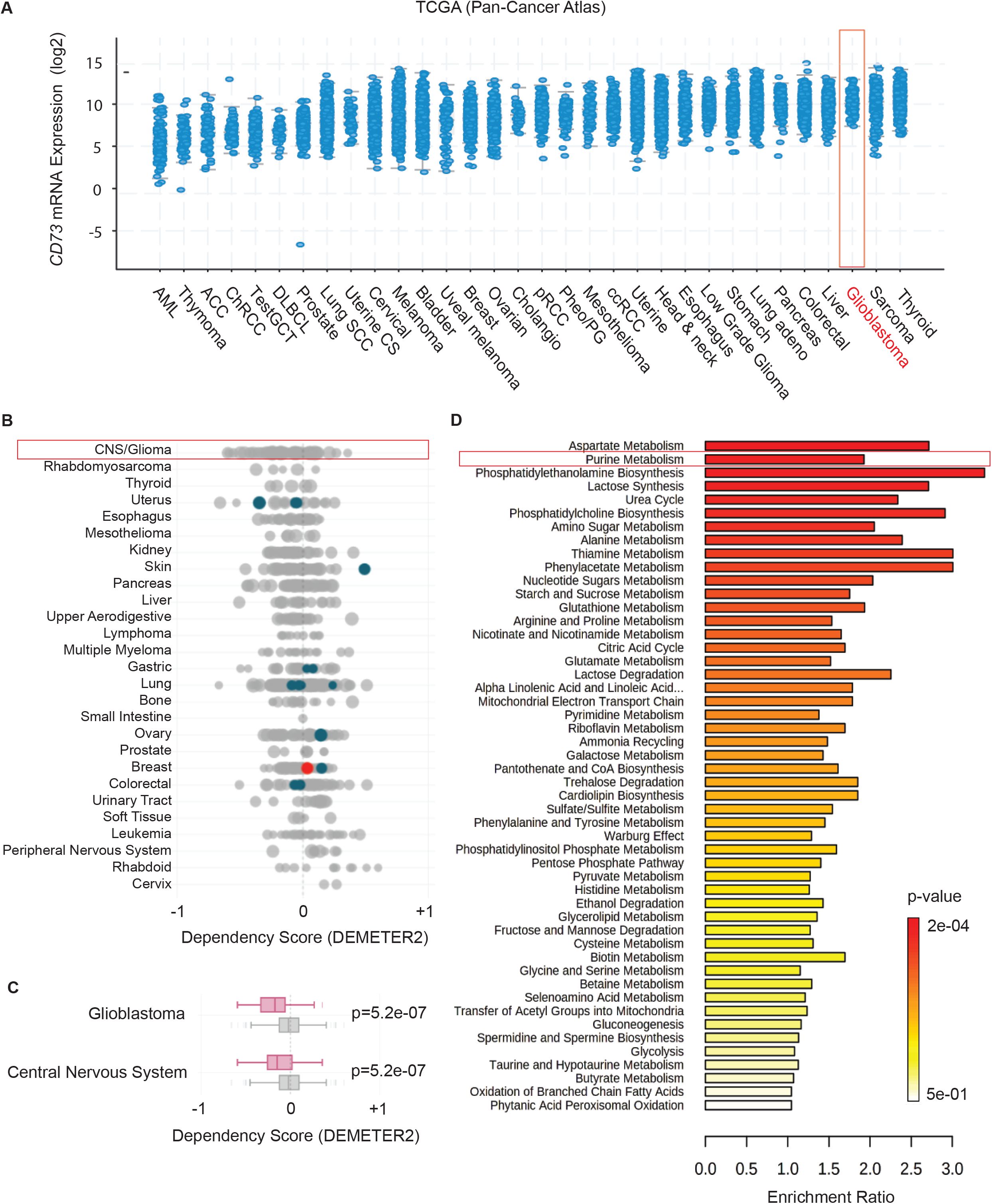
Enrichment of CD73 expression and purine metabolism in glioblastoma and dependency of glioblastoma cell lines on CD73. Bulk *NT5E/CD73* mRNA expression was assessed in tumor tissue (n=10,071) from 32 cancer subtypes (TCGA). Glioblastoma (red box) exhibited high expression of *CD73* relative to most tumor sub-types (**A**). Genome-wide RNAi loss-of-function screening (Project Achilles) in 1,736 cell lines showed that glioblastoma exhibits the highest mean dependence (DEMETER2 score) on *CD73* expression of all cancer lines, while only a small subset of lines from other subtypes depend on *CD73* expression (**B**). Glioblastoma cell lines were the only histologic group which exhibited a significant mean reduction in survival following *CD73* RNAi knockdown (**C**). Mass spectrometry metabolomic profiling (MALDI-MSI) of frozen tissue from 9 IDH-wildtype glioblastoma resections showed significant enrichment of purine metabolism (p<0.05) (**D**).

To determine whether *CD73* expression is intrinsically necessary for tumor cell proliferation and survival, we examined genome-wide RNAi screening data from Project Achilles, a database of gene dependencies for 1,736 cell lines from 34 cancer types^42^. Among all cancers, glioma cell lines exhibited the highest mean dependence on *CD73* expression (**Fig. 1b,c**). Glioblastoma cell lines without isocitrate dehydrogenase-1 and −2 mutations (*IDH1,IDH2*; IDH-wildtype glioblastoma) such as LN340 and U343 exhibited among the highest dependence on *CD73* of all cancer lines, and gliomas were the only lines with a significant mean dependence on *CD73* for survival (p<0.05) (**Fig. 1c**).

To further characterize the significance of purine metabolism in glioblastoma, we used unbiased spatially resolved timsTOF mass spectrometry imaging (MSI) to quantify metabolite levels from multiple pathways. Analysis of ion maps generated from tissue sections from nine fresh-frozen glioblastoma showed that purine metabolism was the second most significantly enriched metabolic pathway. Other enriched pathways included amino acid (aspartate, glutamine, arginine, proline), glutathione and phospholipid metabolism, amino sugar, and nucleotide sugar metabolism (**Fig. 1d**).

### State-specific expression of the core purinergic effectors CD39 and CD73 in glioblastoma by scRNA-seq

Elevated *CD73* expression, significant dependence on *CD73* for cell line survival, and enrichment of purine metabolism prompted us to further explore purinergic signaling in glioblastoma. Given the marked genomic and cellular heterogeneity in glioblastoma^43,44^, we used single-cell analyses to understand which cell types and states are involved in purine pathway activity.

We first analyzed single cell RNA sequencing (scRNA-seq) data from 28 unique human glioblastoma resections encompassing 24,131 total cells^43^. As previously described, most cells in these specimens clustered into four broad lineages according to multi-gene expression signatures, including tumor cells, oligodendrocytes, ‘T cells’, and ‘macrophages’ (**Fig. 2a**). We validated cell classification by quantifying and examining expression of core lineage factors by t-distributed stochastic neighbor embedding (tSNE), and found appropriate restriction for *SOX2* in tumor cells, *PTPRC*/CD45 in immune cells, *SPI1/PU.1, CD68*, and *CD163* in ‘macrophages’, *CD3G* in ‘T cells’, *OLIG1/2* and *MBP* in oligodendrocytes (**Fig. 2a,b; Supplementary Fig. S2a**). In the brain, myeloid markers such as *SPI1/PU. 1, CD68*, and *CD163* may be expressed by tissue-resident microglia or infiltrating peripheral macrophages and dendritic cells; we accordingly classified this category as ‘myeloid’ (see below). While the ‘T cell’ category was predominantly composed of *CD3G* positive T cells (85.1%), it also included sub-populations of NK cells (*NCAM1/CD56+*) (1.1%), and B cells (*MS4A1/CD20+*) (3.2%); we therefore classified this category as ‘lymphoid’ (**Fig. 2a,b**).

**Figure 2:**
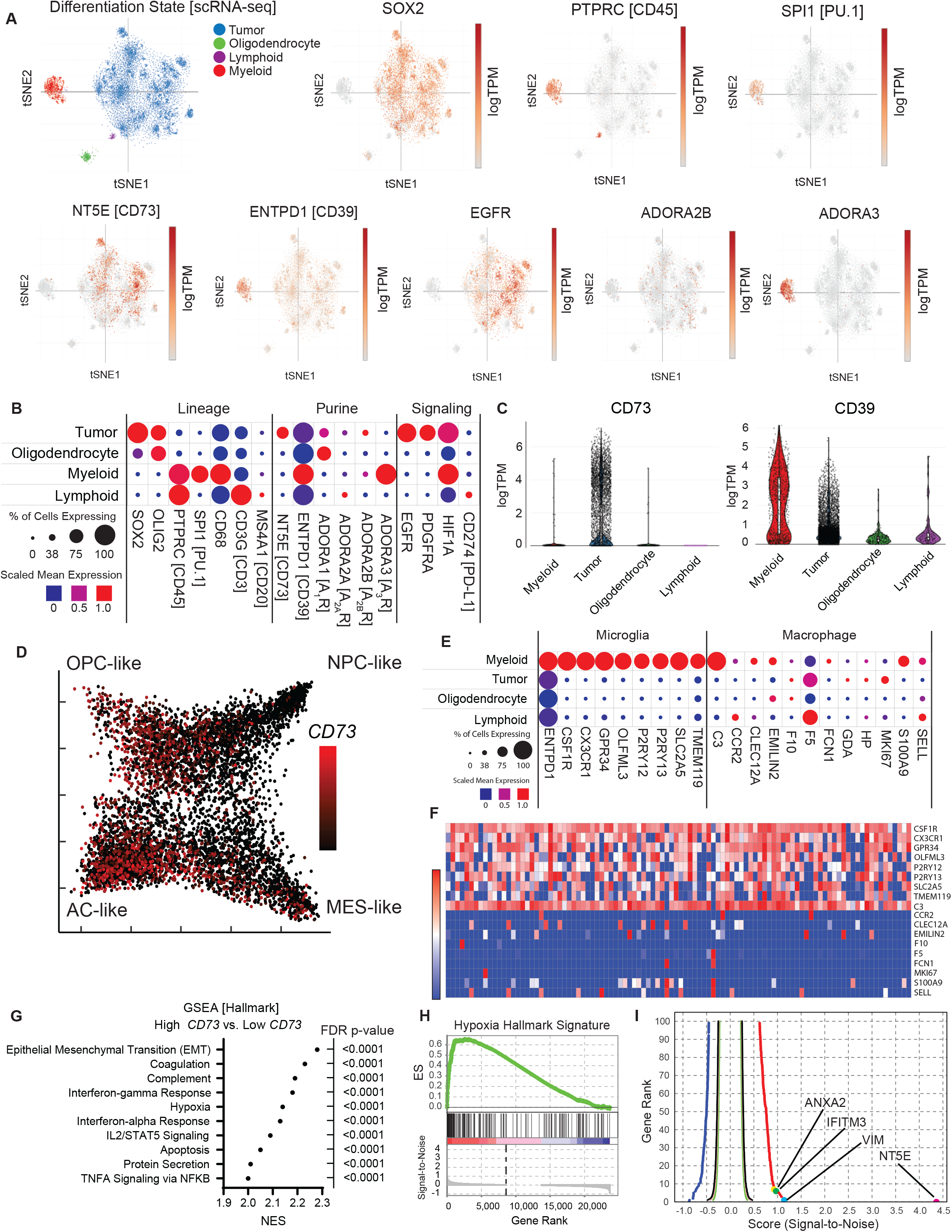
State-specific expression of core purinergic enzymes in glioblastoma. Single-cell RNA-sequencing from 28 human glioblastoma resections encompassing 24,131 cells was analyzed for core purine pathway regulators and related biomarkers. Cells were clustered into lineage sub-groups by expression signature (**A**). Expression patterns were validated by canonical cell type-specific markers, including *SOX2* (tumor), *PTPRC*/CD45 (pan-immune), *SPI1*/PU.1, *CD68* (myeloid), *OLIG2* (oligodendrocyte) and *CD3G/CD3, MS4A1/CD20* (lymphoid) (**A,B**). *CD73* was predominantly expressed by tumor cells (35.9%), with only rare expression in myeloid (2.5%) and other cell types. *CD39* expression was present in all cell types, with higher expression in myeloid cells (98.3% of cells) (**A-C**). Distinct subsets of tumor cells expressed *ADORA1*/A_1_R (24.4%) and ADORA2B/A_2B_R (9.0%), while *ADORA3*/A_3_R was strongly expressed by nearly all myeloid cells (81.8%) (**A,B**). To further characterize the cell states associated with *CD73* expression, tumor cells were clustered according to differentiation state (Neftel *et al.*) into astrocyte-like (AC-like), mesenchymal-like (MES-like), oligodendroglial progenitor cell-like (OPC-like), and neural progenitor cell-like (NPC-like) signatures (**D**). *CD73* expression was strongly enriched in the AC-like signature, which is associated with *EGFR* amplification, with a secondary peak in the OPC-like signature associated with *PDGFRA* amplification. *CD39* expression was correlated with lineage-specific markers of microglia and peripheral macrophages, showing that most myeloid cells in glioblastoma are microglia which exhibit expression of lineage-specific microglial genes such as *TMEM119, P2RY12, CSF1R*, and *CX3CR1* (**E**). Correlation of lineage-markers in cells with the highest levels of *CD39* (top 100 shown) revealed that nearly all exhibited strong microglial expression signatures (**F**). GSEA analysis of tumor cells with the highest (top 1%) and lowest (1%) *CD73* expression showed that pathways including epithelial-to-mesenchymal transition (EMT), interferon (α/γ) response, and hypoxia most strongly associated with strong *CD73* expression (**G,H**), with core genes from each pathway represented among the most enriched genes (**I**).

Prior studies of immune populations isolated from glioblastoma by fluorescence-activated cell sorting (FACS) suggested a role for CD73-expressing macrophages in the pathogenesis of glioma^33^. Analysis of scRNA-seq data confirmed that subsets of myeloid cells do express *CD73*; however, we found that *CD73* was much more prominently expressed by tumor cells (35.9% of cells;1.19 scaled mean expression) than by myeloid cells (2.52%;0.06), or other populations (3.2% oligodendrocytes,0% lymphoid) in this dataset (**Fig. 2a-c**). Conversely, *CD39*, the upstream ectoenzyme that hydrolyzes extracellular ATP and ADP to AMP, while weakly expressed by multiple populations including tumor cells (94.8%;0.73), was more strongly expressed by myeloid cells (98.3%;2.16) (**Fig. 2a-c**).

Analysis of adenosine receptor expression showed that a subset of tumor cells (9.0%) expressed A_2B_R mRNA, a receptor subtype has previously associated with chemoresistance in glioblastoma^45^. A_1_R, which is proposed to inhibit tumor proliferation^46^ was expressed in a distinct subset (24.4%) of predominantly A_2B_R-negative tumor cells at lower levels (**Fig. 2b**). Myeloid cells broadly expressed high levels of A_3_R (81.8% of cells;4.30), while only small subsets expressed other adenosine receptors. In this context, A_3_R is typically associated with microglia and macrophages with immunomodulatory (M2) differentiation^47^. A small subset of lymphoid cells expressed A_2A_R (7.4%), while oligodendrocytes predominantly expressed A_1_R (45.6%).

Single-cell analyses have indicated that glioblastoma cells differentiate into four principle states mirroring developmental stages of the human brain or injury response programs, including astrocyte-like (AC-like), oligodendrocyte progenitor-like (OPC-like), neural progenitor cell-like (NPC-like), and mesenchymal-like (MES-like)^43^. These states are associated with specific underlying genomic alterations, including *EGFR* amplification (AC-like), *PDGFRA* amplification (OPC-like), *CDK4* amplification (NPC-like), and *NF1* loss (MES-like). Analysis of scRNA-seq data showed that tumor cells with AC-like differentiation exhibited the highest *CD73* mRNA expression, while expression was lower in OPC-like and MES-like tumor cells and scarce in the NPC-like cluster (**Fig. 2d**). Analysis of publicly available data in non-neoplastic human (UCSC Cell Browser^48,49^) and mouse (Broad Single Cell Portal^50^) brain showed that *CD73* expression is predominantly enriched in astrocytes and OPC; *CD39* is variably expressed by multiple cell types, but most strongly by microglia and vascular endothelial cells (**Supplementary Figure 2b**).

Myeloid cells, especially microglia, typically constitute the largest fraction of the immune cells in brain tumors^51^. Thus, expression of *CD39* in a large fraction of myeloid cells suggests that microglia may express high levels of *CD39*. In scRNA-seq data^43^, analysis of lineage-specific markers for microglia and peripheral macrophages confirmed this hypothesis, showing that most myeloid cells (70-80%) exhibit an expression signature defined by core microglial genes (*P2RY12, SLC2A5, TMEM119*, and *CX3CR1*) rather than by peripheral macrophage genes (e.g. *CCR2, CLEC12A*, and *F10*), which were present in 20-30% of cells (**Fig. 2e**). *CD39* expression was highly enriched in microglia, which comprise nearly all cells with the highest *CD39* expression (**Fig. 2f**; top 100 *CD39* expressing cells). Single-cell profiling has previously revealed multiple functional microglial categories associated with distinct biological responses^51^. Analysis of scRNA-seq data derived from glioma-associated microglial populations^51^, however, showed no clear correlation between *CD39* or *CD73* expression and specific functional clusters (**Supplementary Fig. S2c**).

To identify gene expression programs associated with *CD73* and *CD39* expression in glioblastoma, we performed gene set enrichment analysis (GSEA)^52^ on scRNA-seq data^43^. CD73 expression in tumor cells was most strongly associated with inflammatory pathways including chemokine secretion (CXCL2), major histocompatibility complex class I (MHC-I) expression, and interferon signaling (**Supplementary Fig. 2d**). Comparing tumor cells with the highest (top 1%) and lowest (bottom 1%) *CD73* expression showed significant enrichment of epithelial-mesenchymal transition (EMT), hypoxia, and interferon alpha and gamma gene sets, along with inflammatory pathways including interferon signaling **(Fig. 2g,h, Supplementary Fig. 2d)**. *CD73* was the most enriched gene in this analysis (signal-to-noise=4.3), with core effectors of each pathway including *VIM* (1.1)(EMT), *IFITM3* (0.9)(IFN-γ), and *ANXA2* (0.9)(hypoxia) among the most enriched genes (**Fig. 2i**). In myeloid cells, *CD39* expression was strongly associated with purine receptor signaling (p=8e-06) along with MHC-II expression and interferon signaling (**Supplementary Fig. S2e**). Collectively, these results suggested that coordinated crosstalk between tumor and immune populations governs immunosuppressive purine metabolism in glioblastoma.

### CD73 protein exhibits distinctive expression patterns in glioblastoma and other central nervous system tumors

To confirm our results of increased CD73 expression in glioblastoma, we next systematically characterized CD73 protein expression in human glioblastomas and other brain tumors by immunohistochemistry (IHC) (**Fig. 3a**). We assessed membranous CD73 protein expression in 566 clinical samples of different histologic and molecular subtypes including gliomas, meningiomas, ependymomas, medulloblastomas, and craniopharyngiomas (**Fig. 3b,c; Supplementary Table 1**). Moderate-to-strong CD73 expression was present in both IDH-wildtype (s.d.±mean;2.2±0.8;n=116) and IDH-mutant glioblastoma (1.9±1.1;n=8;p=0.19), as well as oligodendrogliomas, pilocytic astrocytomas, meningiomas, and papillary craniopharyngiomas (**Fig. 3b,c; Supplementary Fig. S3** and **Supplementary Table 3**). There was slightly reduced CD73 expression in recurrent/residual IDH-wildtype glioblastomas, but not other tumor types (**Supplementary Table 5**). In adamantinomatous craniopharyngioma (ACP), tumor cells were CD73 negative except along cyst lining epithelium, similar to upregulation of PD-L1 by inflammatory mediators in the cyst fluid^53,54^. Peri-tumoral fibrovascular cells also showed widespread CD73 expression in ACP, suggesting that some tumors may exhibit regional or stromal CD73 expression (**Supplementary Fig. S3c** and **Supplementary Table 4**).

**Figure 3:**
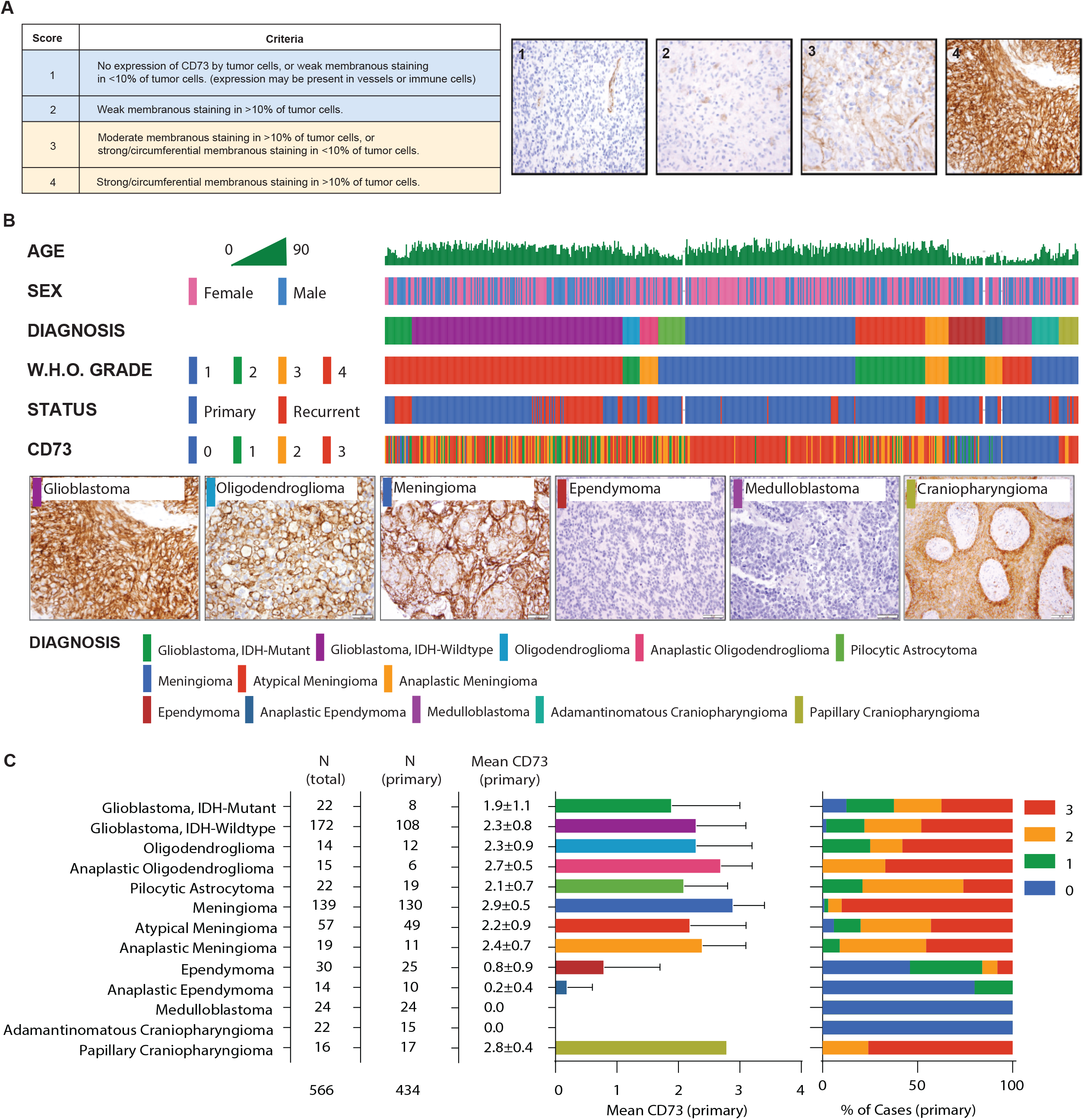
Systematic profiling of CNS tumors demonstrates elevated CD73 expression in glioblastoma. Semi-quantitative immunohistochemistry (IHC) index scoring was developed to characterize CD73 expression in a wide variety of human tissue resections of central nervous system tumors (**A**). IHC was performed on 566 CNS tumor tissue sections, including each major histologic subgroup (**B)**. IDH-mutant [n=22] and IDH-wildtype [n=172] glioblastomas, pilocytic astrocytomas [n=22], and oligodendrogliomas [n=29] frequently exhibited strong CD73 expression. Medulloblastomas of all histologic and molecular subtypes [n=24] and ependymomas [n=44], exhibited minimal to no CD73 expression. Adamantinomatous craniopharyngiomas exhibited no CD73 expression in the tumor epithelium but consistently strong expression in peri-tumoral fibrovascular stroma [n=22], while papillary craniopharyngiomas exhibited strong epithelial CD73 expression in all cases [n=16] (**C**). Scalebars 50μm (B)

### CD73 expression correlates with *EGFR* alteration, clinical outcome, and adenosine level in glioblastoma

From the 194 glioblastomas in our cohort, we analyzed a sub-group of 58 primary IDH-wildtype tumors for clinical or molecular correlations with CD73 expression (**Fig. 4a**). During clinical workup, each case was analyzed using a 447 gene sequencing assay, chromosomal copynumber analysis, and *MGMT* promoter analysis^55^. Polysomy of chromosome 7 and *EGFR* amplification on 7p are highly recurrent events in IDH-wild-type glioblastoma and were present in 66% and 48% of tumors, respectively. Because our scRNA-seq analysis identified a strong correlation between *CD73* expression and an AC-like signature^43^, we explored whether CD73 protein expression was associated with *EGFR* amplification or other genomic alterations in this cohort. Correlative analysis for all common sequence and chromosomal variants was notable only for significantly greater CD73 protein expression (by IHC) in *EGFR* amplified tumors (p=0.029) (**Fig. 4b**).

**Figure 4:**
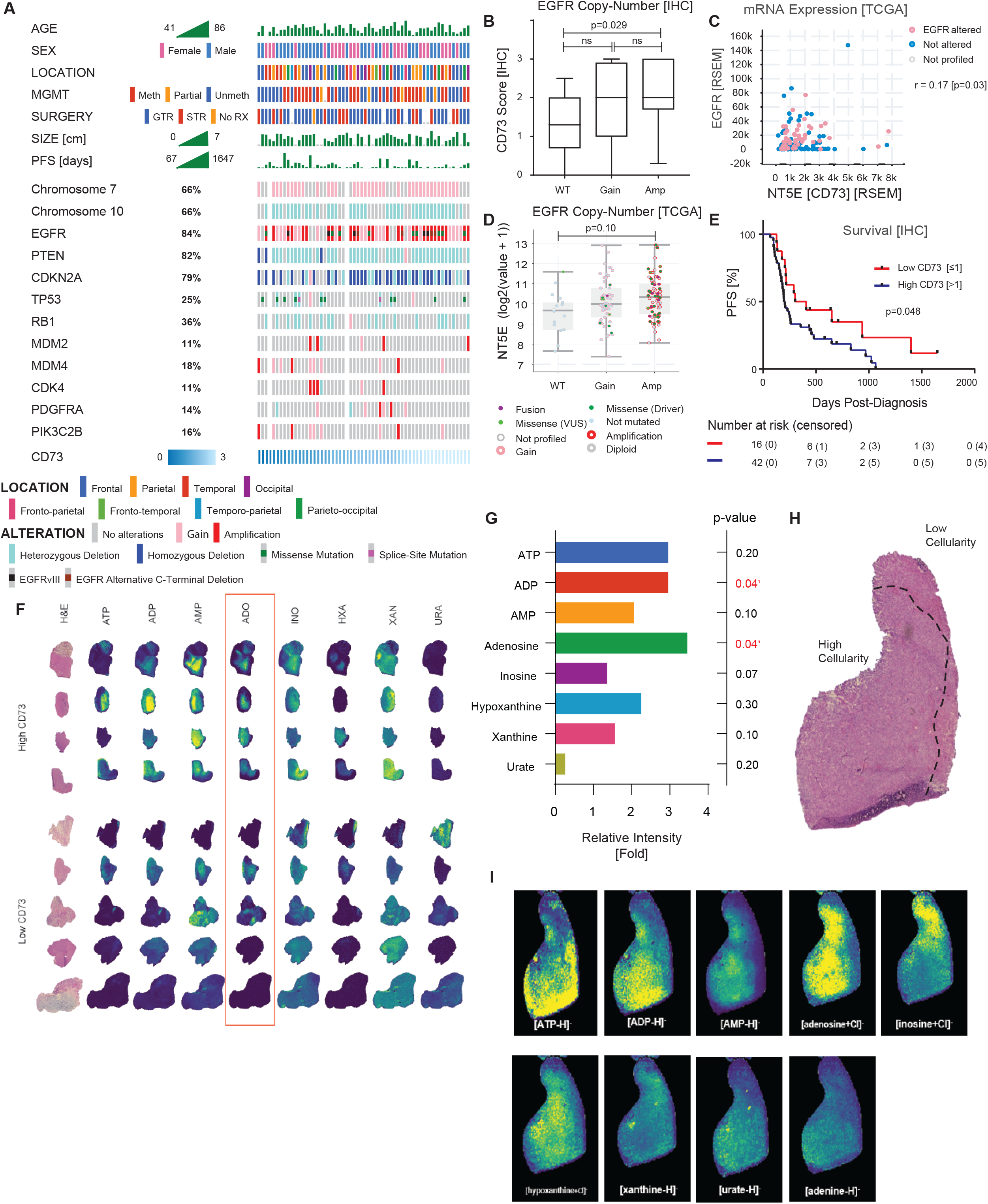
CD73 Expression correlates with genotype and clinical outcome in glioblastoma. To explore the genetic and clinical features associated with CD73 expression in glioblastoma, we evaluated a cohort of 58 primary IDH-wildtype human glioblastoma resections with associated genome-wide chromosomal copy-number (Oncoscan), targeted exome (Oncopanel), and clinical data (**A**). Correlation of CD73 and a wide variety of recurrent genotypic alterations was notable for a significant association between *EGFR* gene amplification and CD73 protein expression (**B**). Analysis of bulk mRNA expression in a broader cohort of IDH-wildtype glioblastoma cases (TCGA) validated these findings, showing a significant direct correlation between *CD73* and *EGFR* expression (**C**). Cases with *EGFR* amplification showed a trend toward elevated *CD73* expression, but this did not reach statistical significance in this dataset (p=0.10) (**D**). Mantel-Cox Log-Rank survival analysis showed that elevated CD73 protein expression (by IHC) was associated with significantly shorter progression-free survival (PFS) (**E**). CD73 IHC was performed on 9 IDH-wildtype glioblastomas previously submitted for mass spectrometry (MSI) showed that 4 cases exhibited high and 5 cases with low CD73 expression. Analysis of purine metabolites in these cases revealed that those with elevated CD73 exhibited significantly higher levels of adenosine relative to cases with low expression (3.5-fold; p=0.04), and it was the most enriched purine metabolite in the tumor microenvironment in CD73-high tumors (**F, G**). Tissue was histologically examined to label regions with high and low tumor cellularity (**H**). Regions of high cellularity exhibit elevated levels of ADP, AMP, and adenosine, while adjacent infiltrative regions with lower cellularity exhibit higher levels of the adenosine catabolic product inosine (**I**) Scalebars 50μm (A). H&E (hematoxylin & eosin), ATP (adenosine triphosphate), ADP (adenosine diphosphate), AMP (adenosine monophosphate), ADO (adenosine), INO (inosine), HXA (hypoxanthine), XAN (xanthine), URA (urate).

To validate these results, we analyzed bulk mRNA sequencing data from TCGA and found a significant positive correlation between *EGFR* and *CD73* mRNA (**Fig. 4c**; r=0.17; p=0.03). There was a strong trend toward increased *NT5E* mRNA expression in *EGFR* amplified tumors, but this association did not reach statistical significance in this dataset (p=0.1) (**Fig. 4d**). There was no correlation between *CD73* expression and *CDK4, PDGFRA*, or *NF1* copy-number alterations or mRNA expression (**Supplementary Fig. S4a-f**). In our cohort, *CD73* expression was slightly lower in recurrent/residual tumors compared to primary lesions (**Supplementary Fig. S4g; Supplementary Table 5**) and in IDH-wild-type glioblastoma with an unmethylated O6-methylguanine-DNA methyltransferase (*MGMT*) promoter compared to *MGMT* methylated tumors (**Supplementary Fig. S4h**). There was no association between *CD73* expression and hypermutation (>10 mutations/MB) or mismatch repair (MMR)-deficiency in recurrent/residual tumors (**Supplementary Fig. S4i**).

Prior studies have shown reduced progression-free survival in glioblastomas with high *CD73* mRNA expression (TCGA)^39,56^. However, patient outcome has not been correlated with tumor cell-specific CD73 protein expression, a more readily applicable measurement in standard clinical practice and clinical trial enrollment. Survival analysis (Mantel-Cox) using semi-quantitative IHC showed significantly longer progression-free survival in IDH-wild-type glioblastoma patients (n=58) with low CD73 protein expression (≤1) compared to those with high CD73 (≥2) (**Fig. 4e**).

We performed CD73 IHC on IDH-wild-type glioblastoma samples that demonstrated enrichment of purine metabolism (**Fig. 1d**)^57^ and found that 4 cases exhibited high CD73 and 5 cases low CD73 expression (**Fig. 4f**). Comparison of whole-tissue metabolite levels from these samples showed that adenosine was the most highly enriched purine metabolite in tumors with high CD73 expression relative to those with low CD73 expression (3.5-fold,p=0.04) (**Fig. 4g**) directly supporting a link between tumor CD73 expression and increased adenosine in glioblastoma tissues. We next registered ion maps with images from adjacent H&E stained tissue sections, and found that ATP, ADP and adenosine were more abundant in cellular tumor regions while inosine, formed by catabolic deamination of adenosine, was enriched in the adjacent infiltrating tumor margin, suggesting regional variations in purine metabolism correlating with tumor density (**Figure 4h, 4i**).

### Multiplexed single cell analysis of glioblastoma tissue

Given cell-state specific expression of critical purinergic regulators such as CD73 and CD39, we next sought to explore the expression, spatial organization, and cell-cell interactions of these proteins in native tissue at sub-cellular resolution. Unlike dissociated scRNA-seq, multiplexed tissue imaging preserves spatial relationships between cells and proteins, enabling correlation of molecular states and tissue architecture^13^,^15^.

We performed 36-marker tissue-based cyclic immunofluorescence (t-CyCIF)^58,59^ on a tissue microarray (TMA) including 139 glioblastoma specimens with associated clinical and genomic data (**Fig. 5a** and **Supplementary Fig. 5**). We segmented individual cells using UNMICST^60^ and ImageJ, and quantified fluorescence intensities on a per-cell basis generating marker expression data from 940,891 cells. Gaussian Mixture Modeling (GMM) was used to assign cell types, and dimensionality reduction of the single cell data was visualized using t-distributed stochastic neighbor embedding (t-SNE) (**Fig. 5b**). Distinct cell-states were identified and analysis of lineage-specific marker expression identified clusters corresponding to distinct classes of cells, including tumor and glia (SOX2, OLIG2, GFAP), lymphoid (CD45, CD3, CD8, CD4), myeloid (CD45, PU.1, CD163, CD63, CD11b), and endothelial cells (CD31) (**Fig. 5b**).

**Figure 5:**
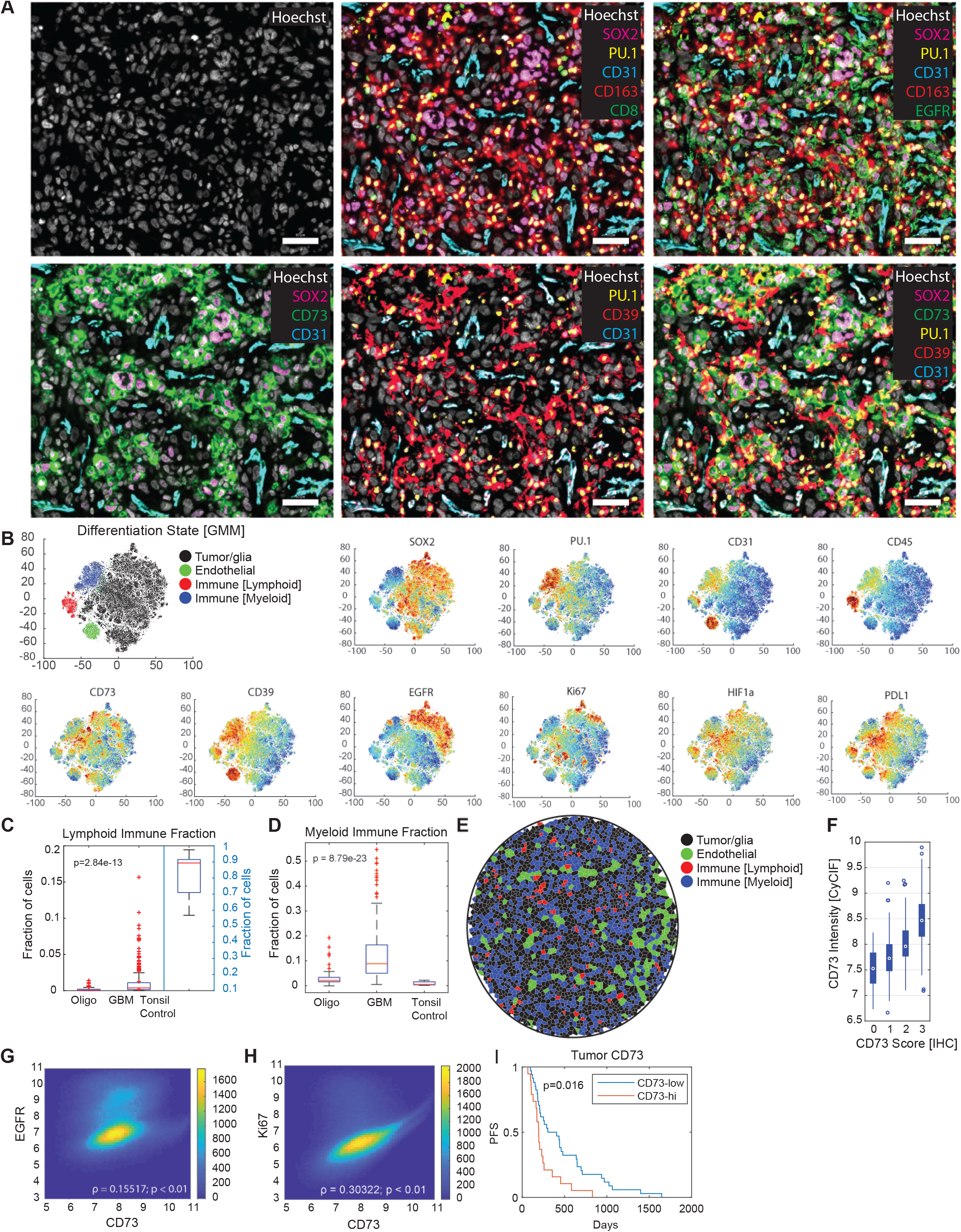
Multiplexed tissue imaging demonstrates state and cell type specific expression of CD73 and CD39 in glioblastoma. 36-plex tissue-based cyclic immunofluorescence (t-CyCIF) encompassing core lineage, signaling, immune, and purine pathway components was performed on a tissue microarray including 172 IDH-wildtype glioblastoma (**A**). Cells were segmented and fluorescence intensity was quantified for each marker at a single-cell level. Cells were clustered by marker expression in an unbiased fashion using Gaussian mixture modeling (GMM) showing 4 dominant signatures which were validated by evaluation of lineage-specific markers. As with transcriptomic analyses, CD73 was predominantly expressed by tumor cells (SOX2+), while CD39 was predominantly expressed by myeloid (PU.1+) and endothelial (CD31+) cells (**B**). Cell type quantification showed that lymphoid populations such as CD3+ T cells were rare in the tumor microenvironment (<5% of cells), though they were more common in glioblastoma than lower grade oligodendrogliomas (**C**), while myeloid cells comprised a substantially larger but variable fraction of the total cell population (**D**). Voronoi tessellation visualization of cell types highlights the non-random distribution of cells and closely intermingled tumor and myeloid cells (**E**) Mean CD73 levels by single tumor-cell analysis showed excellent correlation with IHC score (**F**). Single-cell analysis showed that EGFR protein expression was strongly correlated with CD73 expression (p<0.01) (**G**). Evaluation of other markers was notable for a direct correlation with MIB-1/Ki-67 (**H**) and HIF1a (**see Supplemental Fig. 6**). Higher mean levels of CD73 by singlecell CyCIF analysis correlated with significantly shorter PFS (p=0.016) (**I**).

Single-cell profiling of immune populations revealed that only a small fraction of cells were lymphoid in IDH-wild-type glioblastoma samples (n=172). Comparison with a cohort of low-grade oligodendrogliomas (n=14), showed an even lower proportions of lymphoid cells compared to glioblastoma (**Fig. 5c**). However, tumors frequently contained a substantial proportion of myeloid cells (**Fig. 5d**), confirming that these cells represent the predominant immune population in glioblastoma. To begin to understand the spatial distribution of immune and other cell types, we performed Voronoi tessellation demonstrating a non-random cell distribution including discrete neighborhoods of tumor and myeloid cells (**Fig. 5e**).

Consistent with our observations from scRNA-seq data, CD73 expression was expressed in a subset of myeloid cells, often at similar intensity levels to tumor cells when present (**Supplementary Fig. 6a**)^33^. However, we observed that CD73 expression was more often associated with tumor cells in glioblastoma samples (**Fig. 5a,b**). Mean CD73 expression in marker-defined tumor cells assessed at single-cell resolution by multiplexed immunofluorescence confirmed tissue-level CD73 expression score by IHC (p<0.05) (**Fig. 5f**). Weak CD39 expression was frequently present on tumor cells, however, quantitative analysis and visual inspection showed that strong expression was generally associated with myeloid and endothelial cells (**Fig. 5a,b; Supplementary Fig. 6b**). EGFR and GFAP were predominantly expressed by tumor cells, often heterogeneously, with higher levels in *EGFR* amplified tumors (**Supplementary Fig 6c-d**), consistent with AC-like differentiation of these tumors.

Single-cell analysis showed a strong positive correlation between EGFR and CD73 expression in individual tumor cells. This correlation was also present in tumor cells from cases lacking activating *EGFR* alterations (**Fig. 5g**). Thus, tumors without bulk evidence of genetic activation of *EGFR* may still harbor focal sub-populations of tumor cells with high levels of EGFR and CD73, which may serve to modulate the local immune microenvironment. Examination of other markers was notable for strong direct correlations between CD73 and the proliferation marker Ki-67 (**Fig. 5h**) and HIF1α (**Supplementary Fig. 6e,f**)^61^. Single-cell analysis showed that cases with high mean tumor cell CD73 expression exhibited significantly shorter progression-free survival (p=0.016) (**Fig. 5i**).

### Spatial patterning of core purine regulators in glioblastoma

Compartmentalization of CD39 and CD73 into myeloid and tumor cells, respectively, suggested that the distribution and the positioning of these cell types may influence purinergic metabolism and signaling, with consequent effects on immune activity and tumorigenesis. To study these spatial interactions, we first acquired 18-plex high-resolution images using a Deltavision Elite microscope (**Fig. 6a**). Following signal deconvolution, we generated 3D reconstructions of selected images allowing for precise mapping of marker expression and co-localization (**Fig. 6b,c**). In multiple samples, we observed CD73-expressing SOX2+ tumor cells clustered in close-proximity with CD39-expressing PU.1+/CD163+ myeloid cells, suggesting that functional crosstalk between tumor and immune cells regulates purine metabolism and signaling in glioblastoma.

**Figure 6:**
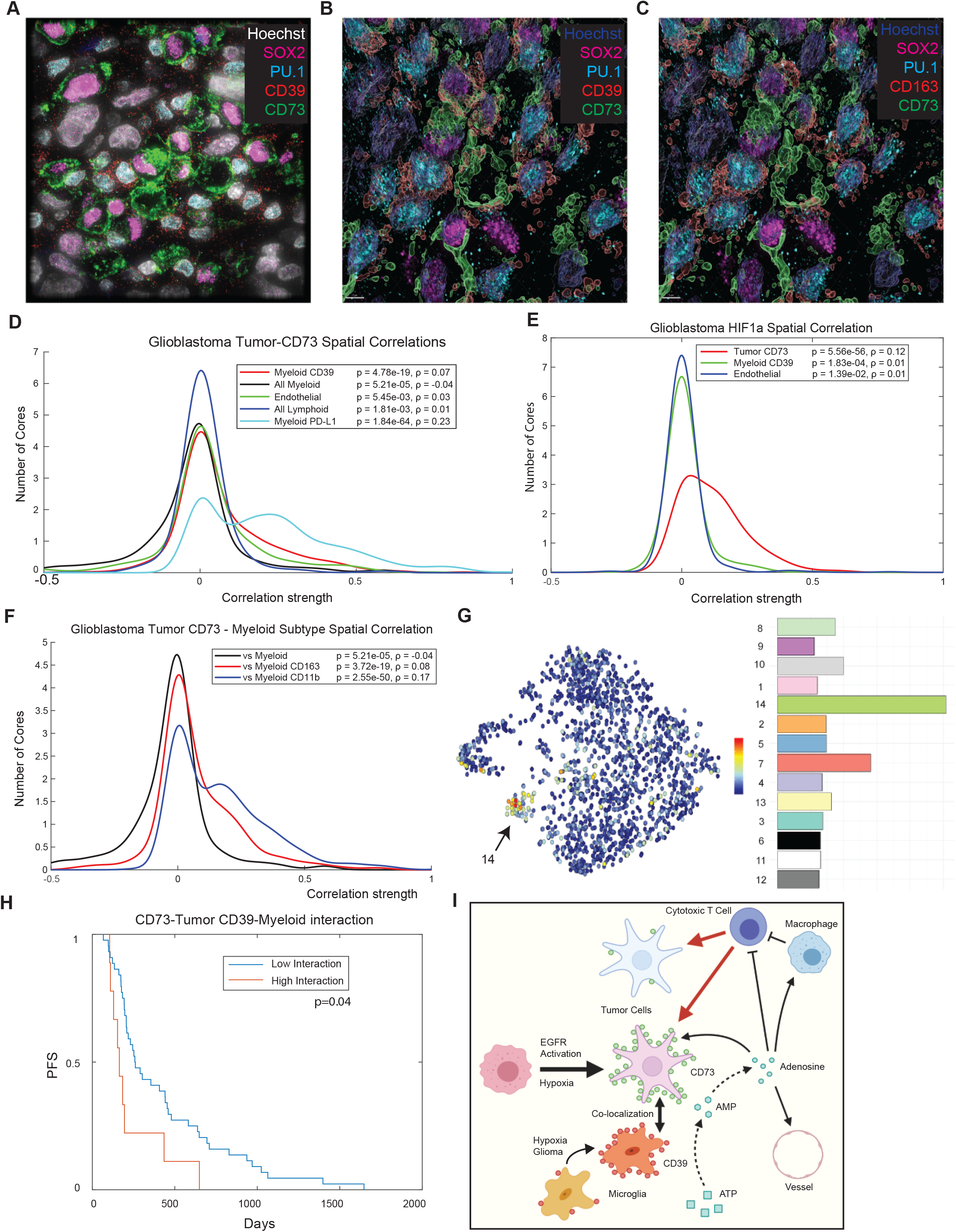
Quantitative single-cell spatial analysis reveals correlation of tumor CD73 and microglial CD39 in glioblastoma and association with clinical outcome. Cyclic multiplexed immunofluorescence with high-resolution deconvolution imaging of glioblastoma specimens was performed using a Deltavision Elite microscope (**A**). 3D rendering of cells and marker expression patterns showed co-localization of tumor CD73 expression and myeloid CD39 expression (**B,C**). To explore this association, spatial neighborhood analysis was performed on the t-CyCIF dataset of 172 IDH-wildtype glioblastoma tissue specimens. This analysis revealed significant spatial correlation between tumor CD73 expression and myeloid CD39 expression (**D**). Tumor CD73 expression does not correlate with the spatial distribution of myeloid cells agnostic for marker expression. Tumor CD73 expression was also significantly correlated with myeloid PD-L1 (**D**) and HIF1a (**E**). Analysis of myeloid differentiation markers showed that tumor CD73 correlated with cells expressing CD163 and CD11b (**F**). Sub-group analysis of microglial cells^51^ showed that these markers correspond to a glioma-associated microglial state (cluster 14) which exhibits elevated inflammatory signaling (**G**). Mantel-Cox survival analysis showed that cases with higher levels of spatial interaction between tumor CD73 and myeloid CD39 showed particularly poor progression-free survival (PFS) (**H**). Collectively, our data suggests a model in which *EGFR* activation and/or hypoxia promotes CD73 expression by glioblastoma tumor cells, and spatial co-localization with CD39-positive microglia leads to elevated adenosine in the tumor microenvironment, promoting immunosuppression and tumorigenesis (**I**).

To assess this association, we used spatial statistics to characterize spatial relationships between tumor, immune, and endothelial cell populations from 748,345 segmented cells from t-CyCIF imaging of 172 IDH-wild-type glioblastomas. We measured the pixel-level correlation strength between CD73 in segmented tumor cells and multiple markers in other populations and compared the spatial distribution of each cell population with respect to underlying marker expression (**Fig. 6d**). CD73 in tumor cells was strongly correlated with CD39 (p=4.78e-19) and PD-L1 (p=1.84e-64) in myeloid cells (**Supplementary Fig. 6e**). There was no convincing spatial correlation between tumor CD73 and myeloid cells themselves agnostic to CD39 or other markers, suggesting that the spatial relationship with CD73 was specific to CD39 and/or PD-L1 expression and not the underlying distribution of cell types. There was no correlation between tumor CD73 expression and endothelial or lymphoid cells. Comparison of tumor CD73 expression and all other t-CyCIF markers was notable for a strong spatial correlation between tumor CD73 and HIF1α present in all cell populations (**Fig. 6e, Supplementary Fig. 6e,6f**).

Because microglia represent most myeloid cells in glioblastoma^62^ (**Fig. 2e**), we examined whether tumor CD73 was spatially co-localized with specific functional microglial classes. We found that tumor CD73 expression was spatially correlated with CD163 and CD11b expression in myeloid cells (**Fig. 6f**). While *CD39* expression was not independently associated with a clear functional microglial signature (**Supplementary Fig. 2b**), combined expression of CD39, CD163, and CD11b revealed strong enrichment for a specific microglial subtype (cluster 14) that is strongly associated with gliomas characterized by enrichment of inflammatory, interferon, and hypoxia-response genes (**Fig. 6g**)^51^. Consistent with this finding, similar pathways were also enriched in our scRNA-seq GSEA analysis of myeloid cells with the highest expression of *CD39* (**Supplementary Fig. 2e**).

If spatial co-localization of tumor CD73 and microglial CD39 increases local production of adenosine by coordinating activity of rate-limiting catabolic enzymes, then tumors with increased spatial correlation may exhibit more aggressive behavior. Survival analysis revealed shorter PFS in tumors with strong spatial interaction of tumor CD73 and myeloid CD39 relative to those with weaker interaction (**Fig. 6h**). While most cases with strong interaction scores (87.5%,n=56) also exhibited strong/diffuse expression of CD73, a subset (12.5%,n=8) had low overall expression of CD73 with clusters of interacting tumor and myeloid cells, with dismal progression-free survival (<200 days) in 3/8 (37.5%) of these cases. Thus, in addition to overall levels of CD73 expression, spatial co-localization of tumor CD73 and myeloid CD39 may promote aggressive behavior and poor clinical outcome even in tumors without diffuse expression of CD73.

Collectively, our findings suggest a model in which tumor cells with aberrant upregulation of CD73 - typically associated with *EGFR* activation, hypoxia, and astrocyte-like differentiation - spatially co-localize with CD39 and CD163 expressing microglia to increase local adenosine levels and drive proliferation, immunomodulation, and poor clinical outcomes (**Fig. 6i**).

## DISCUSSION

In this study we integrate data from several methodologies including single cell RNA-sequencing, spatially resolved mass spectrometry, and highly-multiplexed tissue imaging to study immunomodulatory purinergic signaling in glioblastoma. By enumerating cell types and cell states and analyzing their spatial relationships, we show that components of the purinergic pathway are enriched in distinct tumor and immune cell types and that tumor CD73+ and myeloid CD39+ cells physically co-localize in tumor-immunologic niches, resulting in increased tumor proliferation and microenvironment immunomodulation.

Molecular analyses including single-cell genomic and proteomic approaches can yield important insights into the heterogeneous cell states within tumors. These methods typically employ tissue-dissociative methods which eliminate the spatial associations between tumor cells and other components of the tumor microenvironment thereby preventing the evaluation of tissue-level phenomena which can be critical for promoting tumorigenesis^15–17^. An important recent study using dissociative methodologies identified a distinct population of tumor-infiltrating macrophages that express CD73 in glioblastoma^33^; these cells may persist through recurrence and promote resistance to immunotherapeutic treatment. The authors demonstrated reduced glioblastoma tumor growth in CD73-/- mice with combined inhibition of CTLA-4 and PD-1 checkpoints^33^ suggesting that modulating the extracellular purine pathway may synergize with checkpoint inhibitors. That work provides important pre-clinical evidence supporting a role for immune cell CD73 expression in treatment resistance; however, tumor cell expression of CD73 was not characterized and spatial interactions with CD39 were not explored.

Using both single cell transcriptomic data (n=28) and high-plex imaging of a large cohort of glioblastoma tissues (n=172) we develop a robust spatial description of purinergic metabolism in human glioblastoma. While a subset of myeloid cells expresses CD73, we demonstrate that tumor cells are the predominant source of CD73 in many cases. Alterations of the *CD73* gene locus are rare in glioblastoma and other tumors, and accordingly unlikely to be the cause of CD73 upregulation. CD73 expression is strongly associated with *EGFR* gene activation and astrocyte-like lineage differentiation state in IDH-wild-type glioblastoma. While a smaller subset of CD73 expressing tumor cells exhibited evidence of OPC-like, NP-like, or MES-like differentiation, we found no significant associations between CD73 expression and corresponding *PDGFRA, CDK4*, or *NF1* alterations, nor associations with other recurrent alterations in a cohort of 58 clinically-profiled cases, supporting that cellular states associated with EGFR signaling may be a dominant driver of CD73 expression in glioblastoma.

Gene-set enrichment analysis (GSEA) of scRNA-seq data showed that CD73 expression by tumor cells is also associated with a range of molecular features including epithelial-to-mesenchymal transition (EMT), hypoxia, cytokine, and interferon signaling pathways. Thus, in addition to EGFR signaling, CD73 expression may be promoted by epigenetic processes including inflammation and hypoxia. *EGFR* alterations, particularly amplification of this locus, are recurrent events in IDH-wild-type glioblastoma and tumors frequently develop a hypoxic microenvironment due to pathognomonic vascular abnormalities^63^. These genetic and microenvironmental associations may provide one potential explanation for the particular enrichment of CD73 and purine metabolism in glioblastoma relative to other cancers.

Clinical analyses have previously shown that bulk *CD73* mRNA expression is associated with poor progression-free survival. However, such analyses do not account for the distinct cell populations in tumor tissue, or directly relate mRNA or protein expression of CD73 by tumor cells to clinical outcome. Here, we show that protein expression of CD73 by tumor cells in particular is associated with shorter progression-free survival, as assessed by segmentation and quantification of individual tumor cells using t-CyCIF multiplexed imaging, or by immunohistochemistry (IHC) using a semi-quantitative scoring system. CD73 IHC provides a rapid and applicable test for clinical evaluation by pathologists; incorporation of multiplexed methods into clinical research practice can provide increasing molecular resolution to guide patient enrollment and biomarker assessment in clinical trials^64^.

While CD73 is predominantly expressed by tumor cells in glioblastoma tissue, the upstream enzymatic regulator CD39 of ATP catabolism is primarily expressed by tumor-associated myeloid cells. Single-cell transcriptomics of CD39-expressing myeloid cells show that most are microglia which express core lineage signature genes such as *P2RY12, TMEM119, CX3CR1*, and *CSF1R*, with fewer cells exhibiting a peripheral macrophage signature. Transcriptomic analysis showed that these cells exhibit a pro-inflammatory phenotype characterized by purine signaling, MHC-I/II, and interferon pathway activity. Further sub-classification of CD39-expressing microglia (which display a strong spatial association with CD73-high tumor cells) indicates that they exhibit a pro-inflammatory state highly correlated with hypoxia in glioma tissue^51^. These results suggest that purine metabolism and the equilibrium of purine metabolites is controlled by cell non-autonomous interactions between distinct cell populations.

To address the cellular and molecular heterogeneity of glioblastoma tissue directly, we performed spatial statistical analyses on single cell measurements derived from multiplexed imaging data to correlate expression of cell types and markers in physical space. While tumor CD73 expression was not associated with the distribution of other cell types independent of marker expression, it was highly correlated with microglial CD39 expression, and these cells exhibited a pro-inflammatory glioma-associated phenotype. Tumor CD73 was also associated with HIF1α expression, confirming the findings from single-cell transcriptomic analysis, and again suggesting that hypoxia, in addition to EGFR amplification, may drive local CD73 expression.

The mechanisms underlying the spatial association of tumor CD73 and microglial CD39 are not certain and require further study, though GSEA analysis of CD73-high tumor cells showed enrichment of chemokines including CXCL2 which may recruit myeloid cells in cancer^65^. Prior studies have also shown that glioblastoma tumor cells may upregulate expression of CD39 in macrophages via secretion of molecules such as kynurenine^66^. Thus, spatial associations may be promoted by local microenvironmental conditions (e.g., hypoxia), recruitment of myeloid cells by chemokine secretion, and local promotion of CD39 expression. Demonstrating the potential biological and clinical significance of these findings, we show that purine metabolism is associated with spatial interaction between tumor cells with aberrant CD73 expression and nonneoplastic microglia with CD39 expression, and that this spatial interaction may be linked to poor clinical outcomes.

In tumors with elevated CD73 expression, the enzyme is thought to exert its effects on tumor biology through enzymatic elevation of adenosine concentration in the tumor microenvironment with resultant activation of adenosinergic receptors in an autocrine or paracrine fashion. Previous studies of glioblastoma cell line cultures have demonstrated increased ecto-5’-nucleotidase activity compared to primary astrocyte cell cultures suggesting that CD73 expression may correlate with increased adenosine production in these tumors^56^. In this study, we employed a novel spatially resolved mass spectrometry (MALDI-TOF) imaging (MSI) approach to simultaneously and comprehensively analyze the abundance of metabolites in the purinergic pathway in frozen tissue sections from human glioblastoma resection specimens. Tumors with strong CD73 expression exhibited significantly higher levels of adenosine than those with low CD73 expression (3.5-fold), supporting the hypothesis that CD73 may be enzymatically active in tumors and exert biological activity on tumorigenesis through generation of adenosine in the tumor microenvironment. The homeostatic mechanisms controlling purine metabolism and equilibrium of various metabolites including adenosine are complex, involving additional enzymes beyond CD39 and CD73, and further exploration of the mechanisms underlying purine metabolite levels and their biological significance in a large cohort of human CNS tumors requires further study. However, the detection of a correlation between CD73 expression and adenosine concentration provides a proof-of-principle for the incorporating metabolite profiling of human resection specimens by MSI as a complementary approach to IHC and other analytical methods for detecting therapeutically and clinically meaningful biomarkers for clinical trials or other studies.

In a syngeneic mouse model of glioma, inhibition of A_2A_R, an adenosine receptor expressed on CD8 T cells, provided modest but significant survival benefit but addition of inhibitors of other purinergic signaling effectors including CD73 did not provide synergistic benefits^67^; however, CD73 is not intrinsically expressed in this mouse model system, limiting generalization. Monoclonal antibody CD73 inhibitors are in phase 1-2 clinical trials in advanced solid malignancies (NCT02503774, NCT02754141) and recurrent ovarian cancer (NCT03267589), typically in combination with other checkpoint inhibitors, and have shown promising early results and limited toxicity^68^. Given the high expression of CD73 in glioblastoma and association with clinical outcome, targeting this enzyme with single or combination therapy with checkpoint inhibitors likely represents an attractive therapeutic avenue.

In conclusion, this study uses multiple complementary experimental methodologies (transcriptomics, metabolomic, and high-plex imaging) to demonstrate that purinergic signaling is highly enriched and biologically relevant in IDH-wildtype glioblastoma. Elevated CD73 expression is associated with *EGFR* amplification, astrocyte-like lineage differentiation, inflammatory transcriptional states, and poor clinical outcomes. Spatial statistical analysis of multiplexed immunofluorescence data shows for the first time in human glioblastoma tissue that tumor CD73 and microglial CD39 expression are highly spatially coordinated. These results suggest that purine metabolism and tumor-immune interactions are mediated by population-level interactions between tumor and non-tumor cells, and effectively understanding tissue heterogeneity and immune evasion in glioblastoma likely requires analysis of tissue samples with preserved spatial architecture.

## METHODS

### Tissue Characterization

Formalin fixed and paraffin embedded (FFPE) tissue specimens were retrieved from the archives of Brigham and Women’s Hospital with institutional review board (IRB) approval under DFHCC Protocol #10-417 or waiver of consent protocols. The study is compliant with all relevant ethical regulations regarding research involving human tissue specimens. The Principal Investigator is responsible for ensuring that this project was conducted in compliance with all applicable federal, state and local laws and regulations, institutional policies and requirements of the IRB. All cases were reviewed and classified according to the revised WHO 2016 Classification of Tumours of the Central Nervous System (S.C., S.S.). The characteristics of cases including demographics, genotyping, immunohistochemical, and clinical data are provided (**Supplemental Table 1**).

Specimens included 194 glioblastoma specimens (GBM, W.H.O. grade IV) from 179 distinct patients, including 128 primary and 66 recurrent tumors; 5 diffuse intrinsic pontine gliomas (DIPG; H3K27M-mutant diffuse midline glioma); 14 oligodendroglioma, IDH-mutant, 1p/19q codeleted specimens (OLIG, W.H.O. grade II) from 13 distinct patients, including 12 primary and 2 recurrent tumors; 15 anaplastic oligodendroglioma, IDH-mutant, 1p/19q co-deleted specimens (AOLIG, W.H.O. grade III), including 6 primary and 9 recurrent tumors; 22 pilocytic astrocytomas (PA, W.H.O. grade I) from 22 distinct patients; 139 meningiomas (MEN, W.H.O. grade I) from 139 distinct patients, including 130 primary and 9 recurrent tumors; 57 atypical meningioma specimens (AMEN, W.H.O. grade II) from 57 distinct patients, including 49 primary resections and 8 recurrent tumors; 20 anaplastic meningiomas (W.H.O. grade III), 29 ependymomas (EP, W.H.O. grade II), 15 anaplastic ependymomas (AEP, W.H.O. grade III), 25 medulloblastomas (MED, W.H.O. grade IV), 21 adamantinomatous craniopharyngiomas (ACP, W.H.O. grade I), 14 papillary craniopharyngiomas (PCP, W.H.O. grade I), and 7 solitary fibrous tumors (SFT) of the meninges. All medulloblastomas were classified by histologic subtype per 2016 W.H.O. criteria (S.C., S.S.), and included 15 classic, 3 desmoplastic/nodular, 1 extensive nodularity, and 5 large/cell anaplastic variants. Molecular data including genotype and/or chromosomal copy-number analysis were available for 11/24 (45.8%) medulloblastomas, and all cases were classified according to molecular subtype per 2016 W.H.O. criteria (S.C., S.S.) (cases without molecular data classified as ‘NOS’), including 13 NOS, 5 SHH, 3 Group 3, and 2 Group 4 tumors. Targeted exome sequencing (Oncopanel exhibited IDH1 R132 or IDH2 R172 mutations and whole-arm co-deletion of chromosomes 1p and 19q by chromosomal microarray and/or sequencing. All ependymomas exhibited predominantly classic histologic features. All PCP had *BRAF* V600E mutation by genotype and IHC and lacked *CTNNB1* mutations. 21/22 ACP had *CTNNB1* activation by genotype and/or IHC and lacked *BRAF* V600E mutations by genotype and/or IHC.

### Immunohistochemistry

FFPE sections were de-paraffinized, dehydrated, and endogenous peroxidase activity was blocked. Antigen retrieval was performed in Dako citrate buffer 123°C+/-2, 45 Seconds, at 15+/2 PSI. Slides were incubated with anti-CD73 antibody, (Abcam, EPR6115, rabbit monoclonal clonal) at 1:5000 for 45 minutes or anti-CD39 antibody, (Abcam, EPR20627, rabbit monoclonal clonal) at 1:1000 for 45 minutes, washed, then incubated with Labeled Polymer-HRP anti-rabbit secondary antibody (DakoCytomation, K4011) for 30 minutes. Slides were then incubated with DAKO DAB+ solution for 3-5 minutes and counterstained with hematoxylin. We considered membranous expression of CD73 to represent physiologically relevant (active) expression of the protein. Images were acquired using an Olympus BX41 microscope and an Olympus DP26 digital camera.

### CD73 Expression Profiling

Immunohistochemical expression of CD73 was evaluated using a semi-quantitative scoring system. In brief, cases were scored according to the following system: **Absent (0):** No expression of CD73 by tumor cells, or weak membranous staining in <10% of tumor cells, **Weak (1):** Weak membranous staining in >10% of tumor cells, **Moderate (2):** Moderate membranous staining in >10% of tumor cells, or strong/circumferential membranous staining in <10% of tumor cells, **Strong (3):** Strong/circumferential membranous staining in >10% of tumor cells. Cases with Absent (1) or Weak (2) tumor CD73 expression may be further classified as ‘CD73 Low’, while those with Moderate (2) or Strong (3) expression may be classified as ‘CD73 High’.

### Single-Cell RNA-Seq Analysis

Single-cell analysis was performed using the data from a previously published study (Neftel et al, 2019)^43^. The processed log TPM gene expression matrix, metadata file, and the hierarchy data were all downloaded from the Broad Single Cell portal (https://singlecell.broadinstitute.org/single_cell). The analysis was performed in R using the Seurat package (Butler et al., 2018) and the figures were plotted using the ggplot package. Pathway enrichment analyses were performed using GSEA (Subramanian et al., 2005) and GO Enrichment Analysis (Ashburner et al., 2000; Mi et al., 2019; 10.1093/nar/gkaa1113)

### MALDI tissue preparation

Frozen resection tissue from 9 glioblastomas, including 4 cases with high CD73 and 5 cases with low CD73 by IHC, were selected for MALDI MSI. Tissue was sectioned to 10 μm, thaw mounted onto indium-tin-oxide (ITO) slides, and serial sections were obtained for H&E staining. A high-resolution image of the whole H&E tissues was obtained through image stitching (Zeiss Observer Z.1, Oberkochen, Germany) using a plan-apochromat lens (20×) using an AxioCam MR3 camera. Matrix preparation of 1,5-diaminonaphthalene hydrochloride was prepared to a concentration of 4.3 mg/mL in 4.5/5/0.5 HPLC grade water/ethanol/1 M HCl (v/v/v). The matrix was sprayed using a TM-sprayer (HTX imaging, Carrboro, NC) with parameters of a flow rate (0.09 mL/min), spray nozzle velocity (1200 mm/min), spray nozzle temperature (75°C), nitrogen gas pressure (10 psi), track spacing (2 mm) and a four pass spray.

### MALDI FT-ICR mass spectrometry imaging

The mass spectrometry imaging experiments were conducted using a 9.4 Tesla SolariX XR FT-ICR MS (Bruker Daltonics, Billerica, MA) in negative ion mode. Using a tune mix solution (Agilent Technologies, Santa Clara, CA) the method was calibrated through the electrospray source. The instrumental parameters included a defined pixel step size of 100 μm that covered the m/z range of 58-1200. Each pixel consisted of 200 shots at a laser power of 12% (arbitrary scale) and frequency of 1,000 Hz. To bias the acquisition to improve the capture of the purine metabolites mass range, constant accumulation of selected ions (CASI) mode was used, in which the Q1 was set to m/z 350 with an isolation window of 415. Ion images and mass spectra were viewed and processed using SCiLS Lab software (version 2019c premium, Bruker Daltonics, Billerica, MA), in which the dataset was normalized to the total ion current (TIC). Metabolites were putatively annotated using accurate mass of Δppm <2.5 and cross matched using the Metlin metabolite database.

### t-CyCIF Protocol

Tissue-based cyclic immunofluorescence (t-CyCIF) was performed as described (Lin et al., 2018) and in and at protocols.io (dx.doi.org/10.17504/protocols.io.bjiukkew). In brief, the BOND RX Automated IHC/ISH Stainer was used to bake FFPE slides at 60°C for 30 min, dewax using Bond Dewax solution at 72°C, and perform antigen retrieval using Epitope Retrieval 1 (LeicaTM) solution at 100°C for 20 min. Slides underwent multiple cycles of antibody incubation, imaging, and fluorophore inactivation. All antibodies were incubated overnight at 4°C in the dark. Slides were stained with Hoechst 33342 for 10 min at room temperature in the dark following antibody incubation in every cycle. Coverslips were wet-mounted using 200μL of 10% Glycerol in PBS prior to imaging. Images were taken using a 20x objective (0.75 NA) on a CyteFinder slide scanning fluorescence microscope (RareCyte Inc. Seattle WA). Fluorophores were inactivated by incubating slides in a 4.5% H2O2, 24mM NaOH in PBS solution and placing under an LED light source for 1 hr.

### t-CyCIF image processing and data quantification

Image analysis was performed with the Docker-based NextFlow pipeline MCMICRO (Schapiro et al., 2021) and with customized scripts in Python, ImageJ and MATLAB. All code is available on GitHub. Briefly, after raw images were acquired, the stitching and registration of different tiles and cycles were done with MCMICRO using the ASHLAR module. The assembled OME.TIFF files from each slide were then passed through quantification modules. For background subtraction, a rolling ball algorithm with 50-pixel radius was applied using ImageJ/Fiji. For segmentation and quantification, UNMICST2 was used (Schapiro et al., 2021) supplemented by customized ImageJ scripts (Lin et al., 2018) to generate single-cell data. More details and source code can be found at www.cycif.org.

### Single-cell data QC for t-CyCIF

Single-cell data was passed through several QC steps during generation of cell feature table. Initial QC was done simultaneously with segmentation and quantification, so that cells lost from the specimen in the later cycles would not be included in the output. Next, single-cell data was filtered based on mean Hoechst staining intensity across cycles; cells with coefficient of variation (CV) greater than three standard deviation from the mean were discarded as were any objected identified by segmentation as “cells” but having no DNA intensity. These steps are designed to eliminate cells in which the nuclei are not included as a result of sectioning. Highly auto-fluorescent (AF) cells (measured in cycle 1 or 2) were also removed from the analysis, using a customized MATLAB script that applied a Gaussian Mixture Model (GMM) to identify high-AF population. More details and scripts can be found online (https://github.com/sorgerlab/cycif).

### Spatial Analysis

Spatial correlation functions were evaluated as the Pearson correlation between pairs of cells as a function of their relative distance. Specifically, the Pearson correlation between two variables U and V was evaluated between a cell of type A and its k’th nearest neighbor of type B, and the average distance to k’th nearest neighbors was evaluated in micrometers. In each correlation function A and B were some combination of the 4 GMM classes assigned to each cell (e.g. Tumor and Myeloid, Tumor and All, Tumor and non-Tumor). U and V were either marker log-intensity values or a binary variable indicating whether a cell was or was not of a given type (e.g. CD73 and CD39 values, HIF1a values and Endothelial binary). In Supplementary Fig. S6i, A = Tumor, B = Myeloid, U = CD73, V = CD39.

Spatial correlation functions were evaluated for each core up to k=200, and then fit to a single exponential with two parameters:

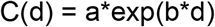

for the correlation C(d) as a function of distance d. The fitted parameter a, correlation strength, was used as a summary of the spatial correlation in each core (Fig. 6d-f). If the 95% confidence interval of a contained 0, a was considered statistically insignificant and set to 0 for downstream analysis. High interaction in Fig. 6h was defined as the condition that all TMA cores from a given patient having significant values for a.

---

**Supplemental Figure 1**

*NT5E/CD73* gene alterations are uncommon in cancer (TCGA), and present in <1% of glioblastomas (**A**). *CD73* mRNA expression levels are not significantly correlated with underlying alteration of the *CD73* gene in glioblastoma (**B**).

**Supplemental Figure 2**

Evaluation of additional lineage-specific markers in single-cell RNA-sequencing data shows expression of *OLIG1, OLIG2*, and *MBP* in oligodendrocyte populations; *CD68, CD163* in myeloid populations; *CD3G* in lymphoid populations. Myeloid cells are enriched for *P2RY12* and *TMEM119,* suggesting that most are microglia rather than infiltrating peripheral macrophages (**A**). Analysis of non-neoplastic human developing cortex (UCSC Cell Browser) showed that *CD73* is predominantly enriched in astrocytes and OPC. *CD39* showed a broader expression distribution but was more strongly expressed by microglia and endothelial cells (**B**) *CD39* and *CD73* expression are not clearly correlated with any specific microglial subtype^51^ (**C**). GSEA analysis of tumor cell populations showed that chemokine (CXCL2), major histocompatibility (MHC-I) and interferon signaling are the pathways most strongly correlated with high *CD73* expression (**D**). GSEA analysis of myeloid cells showed that in addition to purinergic signaling, inflammatory pathways including MHC-II and interferon signaling, and those regulating microglial migration and activation are the most upregulated in cells with high *CD39* expression (**E**).

**Supplemental Figure 3**

Extended immunohistochemistry analysis shows that W.H.O. grade 1 meningiomas exhibit consistently high CD73 expression in all histologic subtypes. Lower CD73 expression is observed in atypical and chordoid meningiomas (W.H.O. grade 2), with the lowest levels in highgrade anaplastic and rhabdoid meningiomas (W.H.O. grade 3) (**A**). Medulloblastomas exhibited no CD73 expression regardless of histologic or molecular subtype (**B**). Adamantinomatous craniopharyngiomas exhibited regional variations in CD73 expression, with no expression in the basaloid, stellate reticular, or whorled tumor epithelium, but consistently strong expression in the fibrovascular stroma, keratinized cyst-lining epithelium, and regions of wet keratin (**C**). Scalebars 50μm (stromal, cystic), 100μm (wet keratin) (C)

**Supplemental Figure 4**

Bulk mRNA analysis of human glioblastoma specimens (TCGA) shows no correlation between *CD73* expression level and chromosomal alterations in *PDGFRA* (**A**), *CDK4* (**B**), or *NF1* (**C**). There was no significant correlation between mRNA expression of *CD73* and *PDGFRA* (**D**), *CDK4* (**E**), or *NF1* (**F**). Mean CD73 protein expression level (by IHC) is slightly reduced in recurrent/residual glioblastomas compared to primary tumors (p<0.05) (**G**). Mean CD73 IHC expression was slightly lower in tumors with an unmethylated *MGMT* promoter, compared to those with methylation of this locus (**H**). There was no significant difference in CD73 IHC expression in recurrent/residual glioblastomas with hypermutation and mismatch-repair deficiency (**I**).

**Supplemental Figure 5**

Tissue-based cyclic immunofluorescence (t-CyCIF) was performed on a tissue microarray (TMA) containing 172 glioblastoma specimens. Representative fields of markers are shown. Scalebars 50μm.

**Supplemental Figure 6**

Quantitative single-cell analysis of CD73 expression in t-CyCIF data shows that tumor and myeloid cells exhibit similar intensity levels of CD73 expression when present (**A**). Myeloid cells exhibit higher levels of CD39 compared to tumor cells (**B**). Single-cell analysis shows that tumor cells from cases with underlying *EGFR* amplification exhibit stronger CD73 expression than cells with polysomy 7, isolated gain-of-function mutations in *EGFR*, or a wild-type diploid *EGFR* locus (**C**). Single-cell analysis shows enrichment of cells with high GFAP expression in *EGFR*-amplified tumors (**D**). t-CyCIF analysis showed spatial correlation of tumor CD73 with HIF1α expression in any population (**E**). Single-cell analysis shows that CD73 expression directly correlate with HIF1α expression in glioblastoma specimens (**F**). Spatial correlation analysis of a representative glioblastoma specimen shows spatial correlation between tumor CD73 and myeloid CD39 expression (**G**). Scalebars 50μm (**E**).

## ACKNOWLEDGEMENTS

We thank Dana-Farber/Harvard Cancer Center in Boston, MA, for the use of the Specialized Histopathology Core, which provided tissue processing services. Dana-Farber/Harvard Cancer Center is supported in part by an NCI Cancer Center Support Grant # NIH 5 P30 CA06516. The results shown here are in part based upon data generated by the TCGA Research Network: https://www.cancer.gov/tcga. This work was supported by NIH grants U54-CA225088 (PKS, SS), R01-CA194005 (SS), T32-GM008313 (SW), and T32-GM007748 (SC), by the Ludwig Center at Harvard (PKS, SS) and by the Specialized Histopathology Core Facility of the Dana-Farber/Harvard Cancer Center (P30-CA06516). KLL was supported by R01CA188288, P01CA163205, R01CA219943.

## AUTHOR CONTRIBUTIONS

SC, MT, and SS developed the concept for the study. JRL, RR, CY, and CCR collected image data and performed analysis in collaboration with SC and SW. SAS, MR, and NYRA designed, performed, and analyzed mass spectrometry experiments in collaboration with SC and SS. SC and PK collected and analyzed single cell RNA-sequencing data. JH collected and analyzed clinical and demographic data in collaboration with SC. All authors wrote and/or edited the manuscript. SS, PKS, NYRA, PB, and KLL provided supervision and funding.

