## Supplementary figures and images for "Single Cell Spatial Analysis Reveals the Topology of Immunomodulatory Purinergic Signaling in Glioblastoma"

### Supplemental Figures

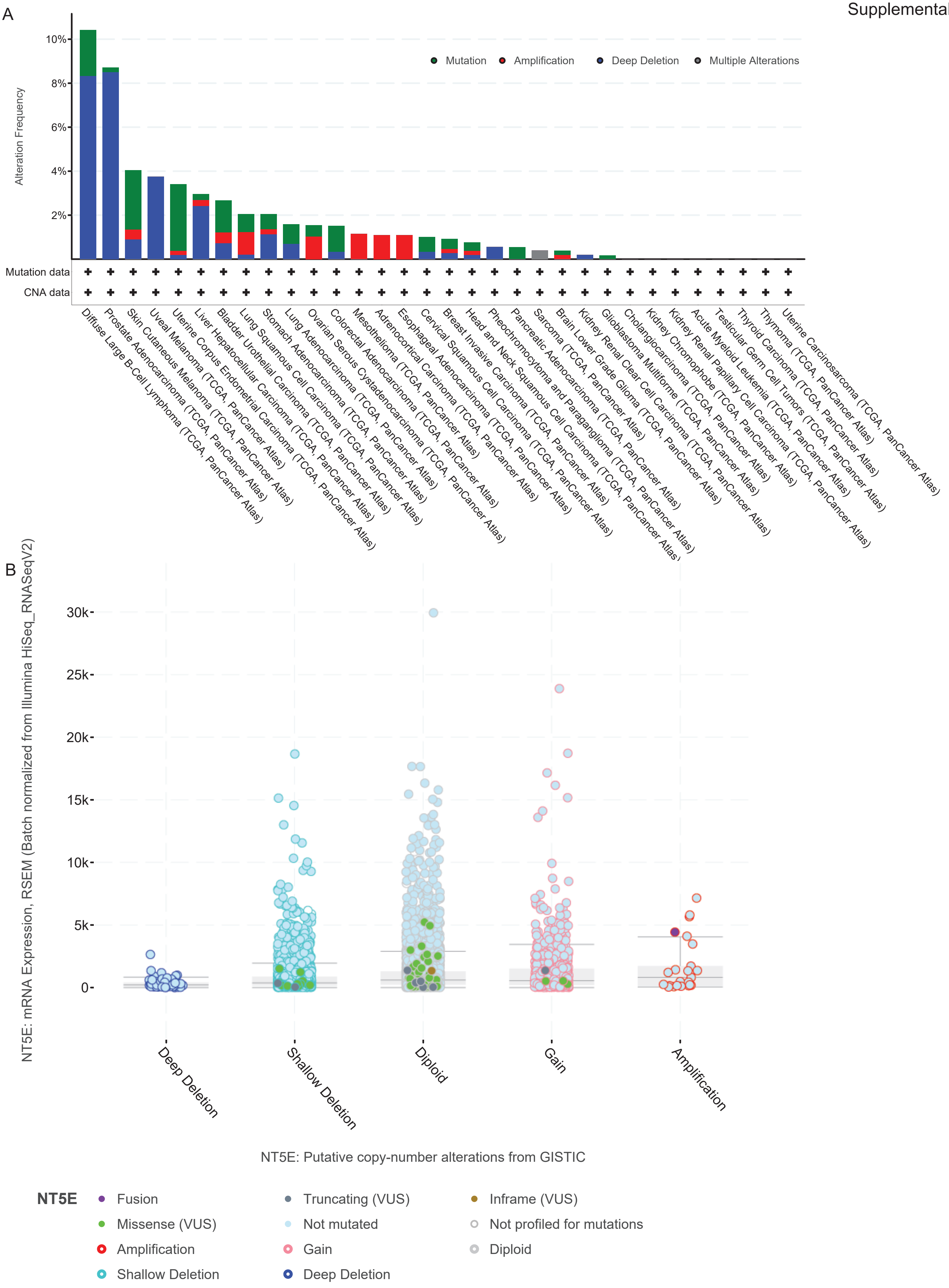

Supplemental Figure 2

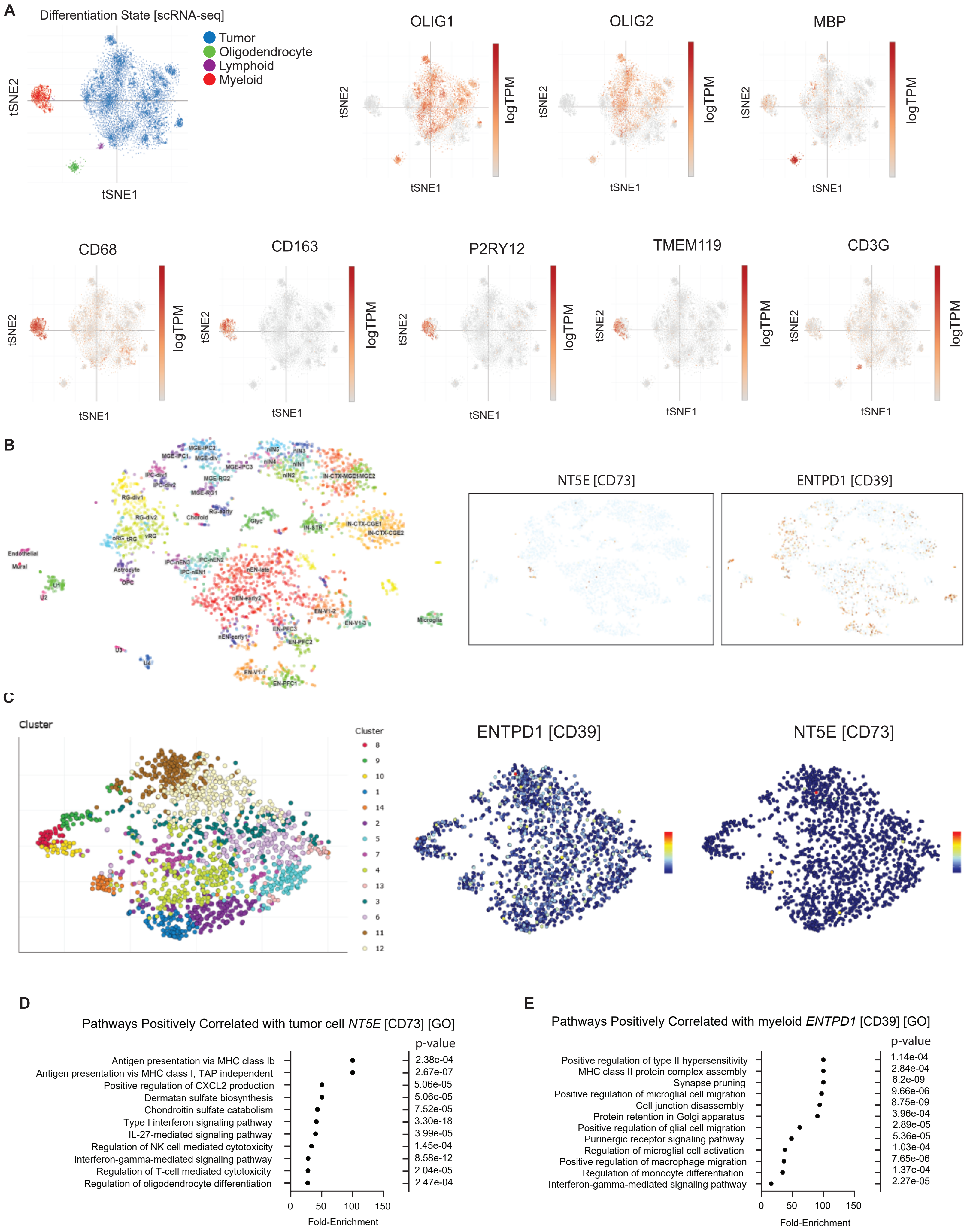

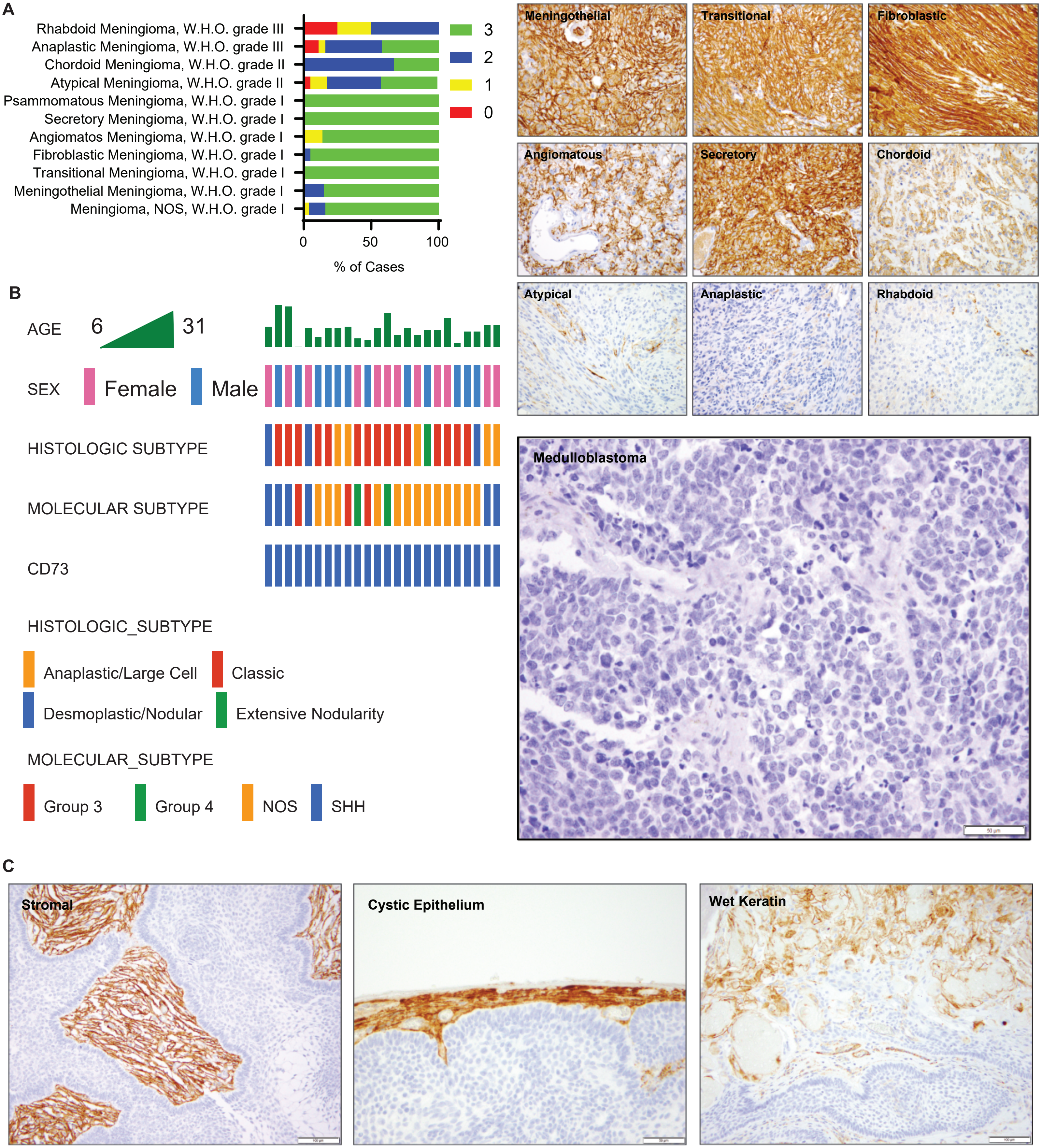

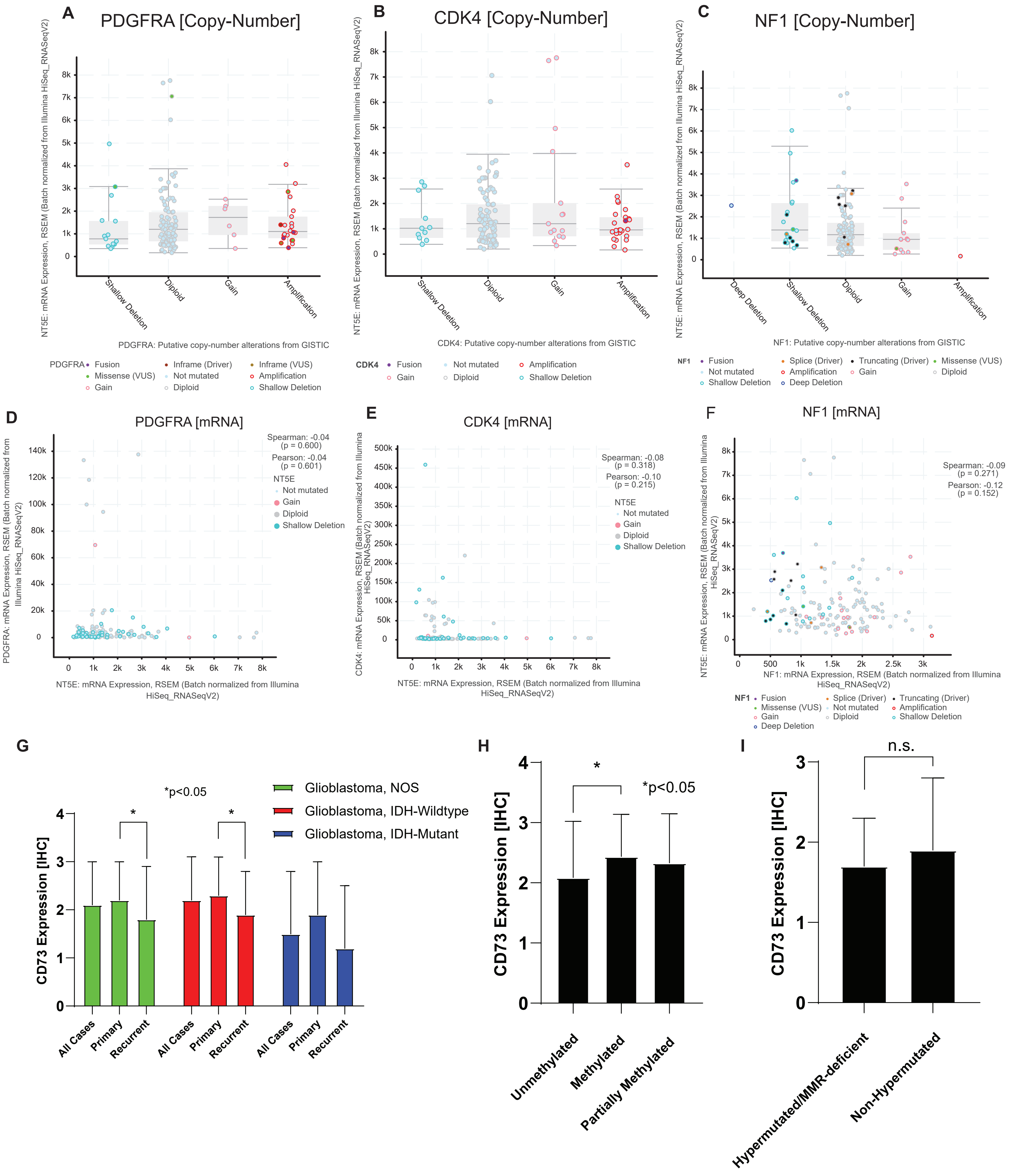

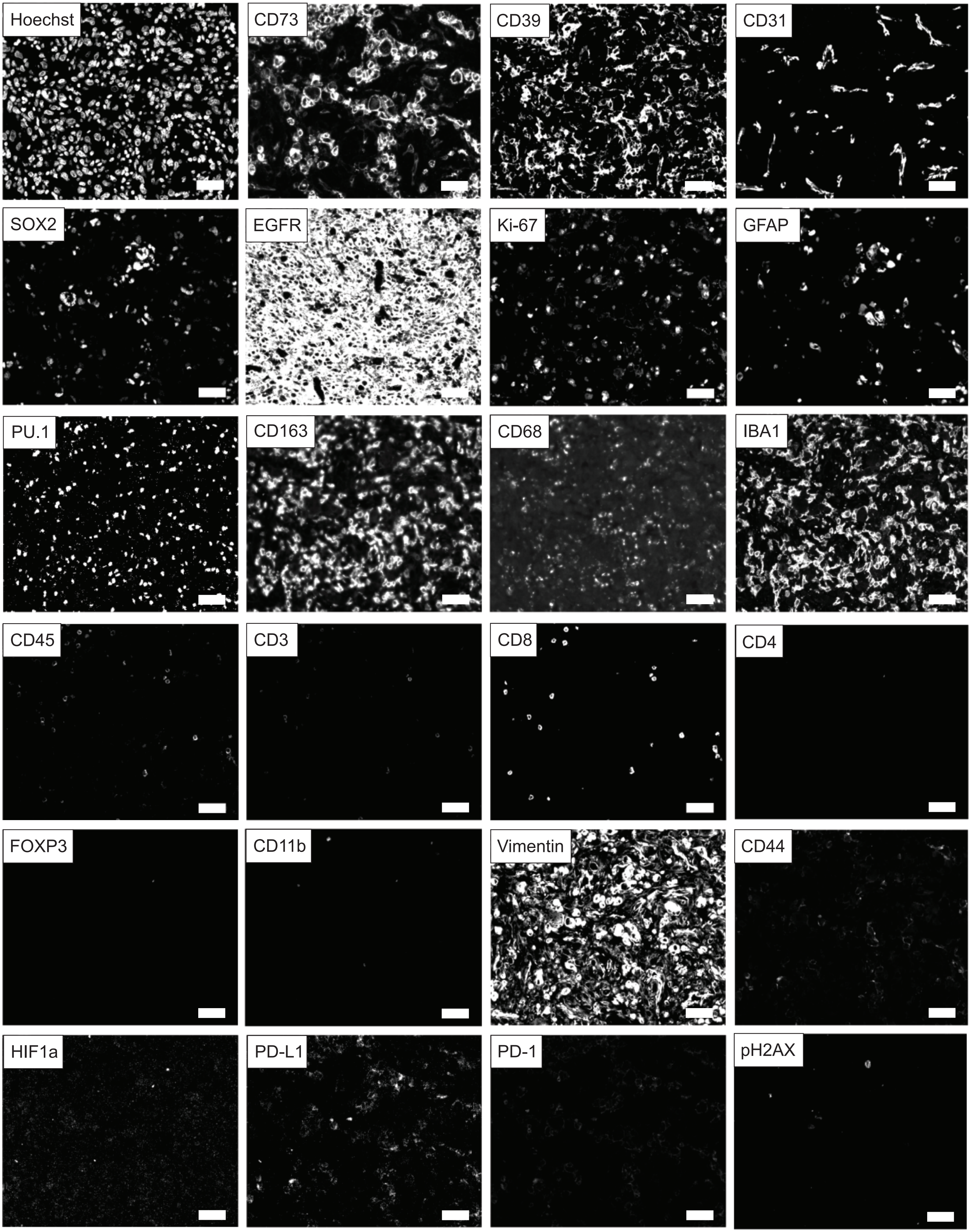

Supplemental Figure 6

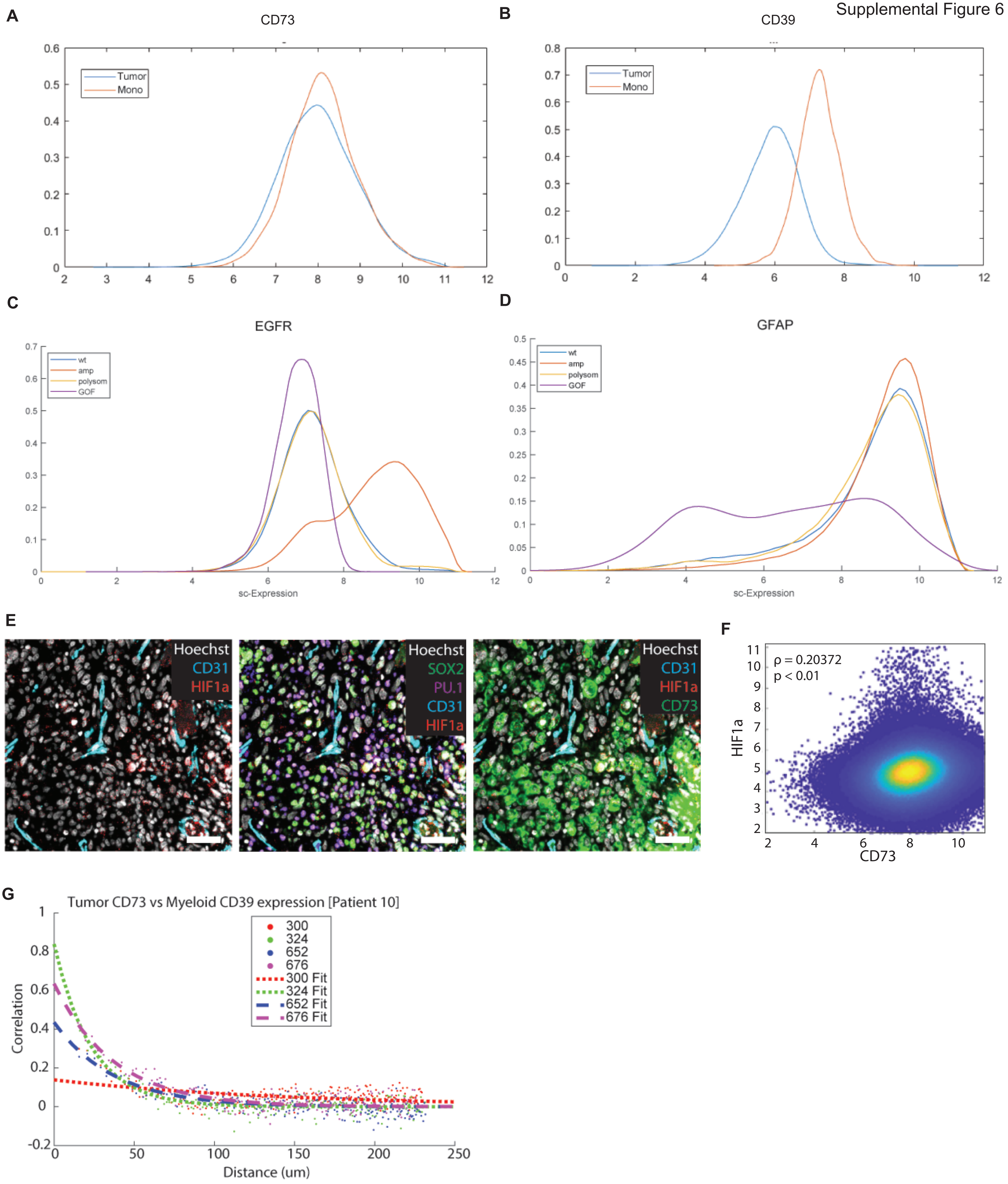
