## Supplemental Tables for "Single Cell Spatial Analysis Reveals the Topology of Immunomodulatory Purinergic Signaling in Glioblastoma"

### Supplemental Table 1: Demographic Data for CNS Neoplasm Cohort

| Diagnosis | W.H.O. | Specimens | Patients | Demographic Data [N (%)] | Age [Mean±S.D.]* | Male [N (%)]ᶎ | Female [N (%)] ᶎ |
|---|---|---|---|---|---|---|---|
| Normal Brain | - | | | | | | |
| Pilocytic Astrocytoma | I | 22 | 22 | 20 (0.91) | 27.9±6.6 | 8 (0.40) | 12 (0.60) |
| Adamantinomatous Craniopharyngioma | I | 22 | 21 | 21 (1.0) | 37.3±17.6 | 12 (0.57) | 9 (0.43) |
| Papillary Craniopharyngioma | I | 17 | 12 | 12 (1.0) | 38.3±15.8 | 5 (0.42) | 7 (0.58) |
| Ependymoma | II | 30 | 29 | 27 (0.93) | 23.1±11.9 | 18 (0.67) | 9 (0.33) |
| Anaplastic Ependymoma | III | 15 | 12 | 11 (0.92) | 28.1±9.4 | 5 (0.45) | 6 (0.55) |
| Medulloblastoma | IV | 24 | 23 | 23 (1.0) | 16.7±6.2 | 12 (0.52) | 12 (0.48) |
| Meningioma | I | 139 | 139 | 139 (1.0) | 57.9±14.0 | 41 (0.29) | 98 (0.71) |
| Atypical Meningioma | II | 57 | 57 | 57 (1.0) | 58.9±14.3 | 27 (0.47) | 30 (0.53) |
| Anaplastic Meningioma | III | 19 | 19 | 19 (1.0) | 59.4±12.2 | 6 (0.32) | 13 (0.68) |
| Oligodendroglioma | II | 14 | 13 | 13 (1.0) | 38.6±12.9 | 5 (0.38) | 8 (0.62) |
| Anaplastic Oligodendroglioma | III | 15 | 15 | 15 (1.0) | 49.5±18.9 | 5 (0.33) | 10 (0.67) |
| Glioblastoma | IV | 194 | 179 | 179 (1.0) | 61.8±11.4 | 68 (0.38) | 111 (0.62) |
| *IDH-Wild Type* | IV | 22 | 22 | 22 (1.0) | 43.3±10.3 | 13 (0.59) | 9 (0.41) |
| *IDH-Mutant* | IV | 156 | 141 | 141 (1.0) | 63.2±10.3 | 76 (0.54) | 65 (0.46) |
| Solitary Fibrous Tumor | - | 7 | 7 | 6 (0.86) | 49.0±19.8 | 1 (0.17) | 5 (0.83) |

*Age of patients at primary diagnosis
ᶎ All patients (primary and recurrent tumors)

### Supplemental Table 2: CD73 Expression in Medulloblastoma by Subtype

| Diagnosis | W.H.O. | N | Score [N (%)] | | | | Mean | S.D. |
|---|---|---|---|---|---|---|---|---|
| | | | 0 | 1 | 2 | 3 | | |
| Medulloblastoma | IV | 24 | 24 (1.0) | 0 (0.00) | 0 (0.00) | 0 (0.00) | 0.0 | 0.0 |
| **Molecular Subtype** | - | - | - | - | - | - | - | - |
| *NOS* | IV | 13 | 13 (1.0) | 0 (0.00) | 0 (0.00) | 0 (0.00) | 0.0 | 0.0 |
| *SHH* | IV | 6 | 6 (1.0) | 0 (0.00) | 0 (0.00) | 0 (0.00) | 0.0 | 0.0 |
| *Group 3* | IV | 3 | 3 (1.0) | 0 (0.00) | 0 (0.00) | 0 (0.00) | 0.0 | 0.0 |
| *Group 4* | IV | 2 | 2 (1.0) | 0 (0.00) | 0 (0.00) | 0 (0.00) | 0.0 | 0.0 |
| **Histologic Subtype** | - | - | - | - | - | - | - | - |
| *Classic* | IV | 15 | 15 (1.0) | 0 (0.00) | 0 (0.00) | 0 (0.00) | 0.0 | 0.0 |
| *Desmoplastic/Nodular* | IV | 3 | 3 (1.0) | 0 (0.00) | 0 (0.00) | 0 (0.00) | 0.0 | 0.0 |
| *Extensive Nodularity* | IV | 1 | 1 (1.0) | 0 (0.00) | 0 (0.00) | 0 (0.00) | 0.0 | 0.0 |
| *Anaplastic/Large Cell* | IV | 5 | 5 (1.0) | 0 (0.00) | 0 (0.00) | 0 (0.00) | 0.0 | 0.0 |

### Supplemental Table 3: Regional Expression of CD73 in Craniopharyngioma

| Diagnosis | W.H.O. | N | Score [N (%)] | | | | Mean | S.D. | p-value* |
|---|---|---|---|---|---|---|---|---|---|
| | | | 0 | 1 | 2 | 3 | | | |
| **Adamantinomatous Craniopharyngioma** | I | - | - | - | - | - | - | - | - |
| ***Total Cases*** | - | 22 | - | - | - | - | - | - | - |
| Basaloid Epithelium | - | 22 | 22 (1.0) | 0 (0.00) | 0 (0.00) | 0 (0.00) | 0.0 | 0.0 | - |
| Stellate Reticular Epithelium | - | 22 | 22 (1.0) | 0 (0.00) | 0 (0.00) | 0 (0.00) | 0.0 | 0.0 | - |
| Whorled Epithelium | - | 22 | 22 (1.0) | 0 (0.00) | 0 (0.00) | 0 (0.00) | 0.0 | 0.0 | - |
| Peri-tumoral Stroma | - | 22 | 0 (0.00) | 0 (0.00) | 3 (0.14) | 19 (0.86) | 2.9 | 0.4 | - |
| Cyst Lining/Keratinizing Epithelium | - | 18 | 1 (0.06) | 0 (0.00) | 9 (0.50) | 8 (0.44) | 2.3 | 0.8 | - |
| Wet Keratin-Associated Cells | - | 21 | 0 (0.00) | 0 (0.00) | 0 (0.00) | 21 (1.0) | 3.0 | 0.0 | - |
| ***Primary Cases*** | - | 15 | - | - | - | - | - | - | - |
| Basaloid Epithelium | - | 15 | 15 (1.0) | 0 (0.00) | 0 (0.00) | 0 (0.00) | 0.0 | 0.0 | - |
| Stellate Reticular Epithelium | - | 15 | 15 (1.0) | 0 (0.00) | 0 (0.00) | 0 (0.00) | 0.0 | 0.0 | - |
| Whorled Epithelium | - | 15 | 15 (1.0) | 0 (0.00) | 0 (0.00) | 0 (0.00) | 0.0 | 0.0 | - |
| Peri-tumoral Stroma | - | 15 | 0 (0.00) | 0 (0.00) | 1 (0.7) | 14 (0.93) | 2.9 | 0.3 | - |
| Cyst Lining/Keratinizing Epithelium | - | 12 | 0 (0.0) | 0 (0.0) | 5 (0.45) | 6 (0.55) | 2.3 | 0.9 | - |
| Wet Keratin-Associated Cells | - | 15 | 0 (0.00) | 0 (0.00) | 0 (0.00) | 15 (1.0) | 3.0 | 0.0 | - |
| ***Recurrent/Residual Cases*** | - | 7 | - | - | - | - | - | - | - |
| Basaloid Epithelium | - | 7 | 7 (1.0) | 0 (0.00) | 0 (0.00) | 0 (0.00) | 0.0 | 0.0 | - |
| Stellate Reticular Epithelium | - | 7 | 7 (1.0) | 0 (0.00) | 0 (0.00) | 0 (0.00) | 0.0 | 0.0 | - |
| Whorled Epithelium | - | 7 | 7 (1.0) | 0 (0.00) | 0 (0.00) | 0 (0.00) | 0.0 | 0.0 | - |
| Peri-tumoral Stroma | - | 7 | 0 (0.0) | 0 (0.0) | 2 (0.29) | 5 (0.71) | 2.7 | 0.5 | **0.2533** |
| Cyst Lining/Keratinizing Epithelium | - | 6 | 0 (0.0) | 0 (0.0) | 4 (0.67) | 2 (0.33) | 2.3 | 0.5 | **1.0000** |
| Wet Keratin-Associated Cells | - | 6 | 0 (0.00) | 0 (0.00) | 0 (0.00) | 6 (1.0) | 3.0 | 0.0 | - |
| **Papillary Craniopharyngioma** | I | - | - | - | - | - | - | - | - |
| ***Total Cases*** | - | 16 | - | - | - | - | - | - | - |
| Squamous Epithelium | - | 16 | 0 (0.00) | 0 (0.00) | 3 (0.19) | 13 (0.81) | 2.8 | 0.4 | - |
| Peri-tumoral Stroma | - | 16 | 3 (0.18) | 5 (0.31) | 9 (0.50) | 0 (0.00) | 1.3 | 0.8 | - |
| ***Primary Cases*** | - | 9 | - | - | - | - | - | - | - |
| Squamous Epithelium | - | 9 | 0 (0.00) | 0 (0.00) | 2 (0.22) | 7 (0.78) | 2.8 | 0.4 | - |
| Peri-tumoral Stroma | - | 9 | 1 (0.11) | 4 (0.44) | 4 (0.44) | 0 (0.0) | 1.3 | 0.7 | - |
| ***Recurrent/Residual Cases*** | - | 7 | - | - | - | - | - | - | - |
| Squamous Epithelium | - | 7 | 0 (0.00) | 0 (0.00) | 1 (0.14) | 6 (0.86) | 2.9 | 0.4 | **0.6275** |
| Peri-tumoral Stroma | - | 7 | 2 (0.29) | 2 (0.29) | 3 (0.43) | 0 (0.0) | 1.3 | 1.0 | **1.0000** |

*p-value relative to primary cases (unpaired t-test)

### Supplemental Table 4: CD73 Expression in Meningioma by Histologic Subtype

| Diagnosis | W.H.O. | N^ζ | Score [N (%)] | | | | Mean | S.D. | p-value* |
|---|---|---|---|---|---|---|---|---|---|
| | | | 0 | 1 | 2 | 3 | | | |
| Meningioma | I | 139 | 1 (0.01) | 3 (0.02) | 11 (0.08) | 124 (0.89) | 2.9 | 0.5 | - |
| *NOS* | I | 69 | 1 (0.01) | 2 (0.03) | 8 (0.12) | 58 (0.84) | 2.8 | 0.6 | 0.2059 |
| *Meningothelial* | I | 13 | 0 (0.00) | 0 (0.00) | 2 (0.15) | 11 (0.85) | 2.8 | 0.4 | 0.4852 |
| *Transitional* | I | 17 | 0 (0.00) | 0 (0.00) | 0 (0.00) | 17 (1.0) | 3.0 | 0.0 | 0.4122 |
| *Fibroblastic* | I | 22 | 0 (0.00) | 0 (0.00) | 1 (0.05) | 21 (0.95) | 3.0 | 0.2 | 0.3567 |
| *Angiomatous* | I | 7 | 0 (0.00) | 1 (0.14) | 0 (0.00) | 6 (0.86) | 2.7 | 0.8 | 0.3187 |
| *Secretory* | I | 9 | 0 (0.00) | 0 (0.00) | 0 (0.00) | 9 (1.0) | 3.0 | 0.0 | 0.5507 |
| *Psammomatous* | I | 2 | 0 (0.00) | 0 (0.00) | 0 (0.00) | 2 (1.0) | 3.0 | 0.0 | 0.7785 |
| Atypical Meningioma | II | 57 | 3 (0.05) | 7 (0.12) | 23 (0.40) | 24 (0.42) | 2.2 | 0.9 | **<0.0001** |
| Anaplastic Meningioma | III | 19 | 2 (0.11) | 1 (0.05) | 8 (0.42) | 8 (0.42) | 2.2 | 1.0 | **<0.0001** |
| Chordoid Meningioma | II | 3 | 0 (0.00) | 0 (0.00) | 2 (0.67) | 1 (0.33) | 2.3 | 0.6 | **0.0422** |
| Rhabdoid Meningioma | III | 4 | 1 (0.25) | 1 (0.25) | 2 (0.50) | 0 (0.00) | 1.3 | 1.0 | **<0.0001** |

ζ Including all cases in cohort
*Relative to W.H.O. grade I meningioma (unpaired t-test)

### Supplemental Table 5: CD73 Expression in Primary and Recurrent Cases

| Diagnosis | W.H.O. | N^ζ | Score [N (%)] | | | | Mean | S.D. | p-value* |
|---|---|---|---|---|---|---|---|---|---|
| | | | 0 | 1 | 2 | 3 | | | |
| **Pilocytic Astrocytoma** | I | - | - | - | - | - | - | - | - |
| *All Cases* | - | 22 | 0 (0.00) | 4 (0.21) | 10 (0.53) | 5 (0.26) | 2.0 | 0.7 | - |
| *Primary* | - | 19 | 0 (0.00) | 4 (0.21) | 10 (0.53) | 5 (0.26) | 2.1 | 0.7 | - |
| *Recurrent* | - | 1 | 0 (0.00) | 0 (0.00) | 1 (1.0) | 0 (0.00) | 2.0 | 0.0 | - |
| **Ependymoma** | II | - | - | - | - | - | - | - | - |
| *All Cases* | - | 30 | 17 (0.57) | 9 (0.30) | 2 (0.07) | 2 (0.07) | 0.6 | 0.9 | - |
| *Primary* | - | 25 | 12 (0.46) | 9 (0.38) | 2 (0.08) | 2 (0.08) | 0.8 | 0.9 | - |
| *Recurrent* | - | 3 | 3 (1.0) | 0 (0.00) | 0 (0.00) | 0 (0.00) | 0.0 | 0.0 | 0.1420 |
| **Anaplastic Ependymoma** | III | - | - | - | - | - | - | - | - |
| *All Cases* | - | 14 | 11 (0.79) | 2 (0.14) | 1 (0.07) | 0 (0.00) | 0.3 | 0.6 | - |
| *Primary* | - | 10 | 8 (0.8) | 2 (0.2) | 0 (0.00) | 0 (0.00) | 0.2 | 0.4 | - |
| *Recurrent* | - | 3 | 3 (1.0) | 0 (0.00) | 0 (0.00) | 0 (0.00) | 0.0 | 0.0 | 0.4189 |
| **Meningioma** | I | - | - | - | - | - | - | - | - |
| *All Cases* | - | 139 | 1 (0.01) | 3 (0.02) | 11 (0.08) | 124 (0.89) | 2.9 | 0.5 | - |
| *Primary* | - | 130 | 1 (0.01) | 3 (0.02) | 9 (0.07) | 117 (0.90) | 2.9 | 0.5 | - |
| *Recurrent* | - | 9 | 0 (0.00) | 0 (0.00) | 2 (0.22) | 7 (0.78) | 2.8 | 0.4 | 0.5585 |
| **Atypical Meningioma** | II | - | - | - | - | - | - | - | - |
| *All Cases* | - | 57 | 3 (0.05) | 7 (0.12) | 23 (0.40) | 24 (0.42) | 2.2 | 0.9 | - |
| *Primary* | - | 49 | 3 (0.06) | 7 (0.14) | 18 (0.37) | 21 (0.43) | 2.2 | 0.9 | - |
| *Recurrent* | - | 8 | 0 (0.00) | 0 (0.00) | 5 (0.63) | 3 (0.38) | 2.4 | 0.5 | 1.0000 |
| **Anaplastic Meningioma** | III | - | - | - | - | - | - | - | - |
| *All Cases* | - | 19 | 2 (0.11) | 1 (0.05) | 8 (0.42) | 8 (0.42) | 2.2 | 1.0 | - |
| *Primary* | - | 11 | 0 (0.00) | 1 (0.09) | 5 (0.45) | 5 (0.45) | 2.4 | 0.7 | - |
| *Recurrent* | - | 8 | 2 (0.25) | 0 (0.00) | 3 (0.38) | 3 (0.38) | 1.9 | 1.2 | 0.2675 |
| **Oligodendroglioma** | II | - | - | - | - | - | - | - | - |
| *All Cases* | - | 14 | 0 (0.00) | 3 (0.21) | 3 (0.21) | 8 (0.57) | 2.4 | 0.8 | - |
| *Primary* | - | 12 | 0 (0.00) | 3 (0.25) | 2 (0.17) | 7 (0.38) | 2.3 | 0.9 | - |
| *Recurrent* | - | 2 | 0 (0.00) | 0 (0.00) | 1 (0.50) | 1 (0.50) | 2.5 | 0.7 | 0.7724 |
| **Anaplastic Oligodendroglioma** | III | - | - | - | - | - | - | - | - |
| *All Cases* | - | 15 | 0 (0.00) | 3 (0.20) | 3 (0.20) | 9 (0.60) | 2.4 | 0.8 | - |
| *Primary* | - | 6 | 0 (0.00) | 0 (0.00) | 2 (0.33) | 4 (0.67) | 2.7 | 0.5 | - |
| *Recurrent* | - | 9 | 0 (0.00) | 3 (0.33) | 1 (0.11) | 5 (0.56) | 2.2 | 1.0 | 0.2811 |
| **Glioblastoma, NOS** | IV | - | - | - | - | - | - | - | - |
| *All Cases* | - | 194 | 11 (0.06) | 45 (0.23) | 53 (0.27) | 85 (0.44) | 2.1 | 0.9 | - |
| *Primary* | - | 128 | 3 (0.02) | 25 (0.2) | 38 (0.30) | 62 (0.48) | 2.2 | 0.8 | - |
| *Recurrent* | - | 66 | 8 (0.12) | 20 (0.30) | 15 (0.23) | 23 (0.35) | 1.8 | 1.1 | **0.0043** |
| **Glioblastoma, IDH-Wildtype** | IV | - | - | - | - | - | - | - | - |
| *All Cases* | - | 156 | 4 (0.03) | 37 (0.24) | 46 (0.29) | 69 (0.44) | 2.2 | 0.9 | - |
| *Primary* | - | 108 | 2 (0.02) | 20 (0.19) | 33 (0.31) | 53 (0.49) | 2.3 | 0.8 | - |
| *Recurrent* | - | 48 | 2 (0.04) | 17 (0.35) | 13 (0.27) | 16 (0.33) | 1.9 | 0.9 | **0.0063** |
| **Glioblastoma, IDH-Mutant** | IV | - | - | - | - | - | - | - | - |
| *All Cases* | - | 22 | 7 (0.32) | 5 (0.23) | 3 (0.14) | 7 (0.32) | 1.5 | 1.3 | - |
| *Primary* | - | 8 | 1 (0.13) | 2 (0.25) | 2 (0.25) | 3 (0.38) | 1.9 | 1.1 | - |
| *Recurrent* | - | 14 | 6 (0.43) | 3 (0.21) | 1 (0.07) | 4 (0.29) | 1.2 | 1.3 | **0.2151** |

*p-value relative to primary cases (unpaired t-test)

### Supplemental Table 6: CD73 Expression by Genotype in IDH-Wildtype Glioblastoma

| Diagnosis | N$^{\zeta}$ | Score [N (%)] | | | | Mean | S.D. | p-value* |
|---|---|---|---|---|---|---|---|---|
| | | 0 | 1 | 2 | 3 | | | |
| **Total Cases** | 108 | 2 (0.02) | 20 (0.19) | 33 (0.31) | 53 (0.49) | 2.3 | 0.8 | - |
| ***MGMT* promoter methylation** | 103 | - | - | - | - | - | - | - |
| *Unmethylated* | 47 | 2 (0.04) | 12 (0.26) | 13 (0.28) | 20 (0.43) | 2.09 | 0.93 | - |
| *Methylated* | 50 | 0 (0.00) | 6 (0.12) | 16 (0.32) | 28 (0.56) | 2.44 | 0.7 | **0.0382** |
| *Partially Methylated* | 6 | 0 (0.00) | 1 (0.17) | 2 (0.33) | 3 (0.50) | 2.33 | 0.82 | 0.5499 |

ζ Cases at primary diagnosis
